# mRNA display reveals a class of high-affinity bromodomain-binding motifs that are not found in the human proteome

**DOI:** 10.1101/2023.05.17.541126

**Authors:** Jason K K Low, Karishma Patel, Paul Solomon, Alexander Norman, Petr Pachl, Natasha Jones, Richard J Payne, Toby Passioura, Hiroaki Suga, Louise J Walport, Joel P Mackay

## Abstract

Bromodomains regulate gene expression by recognizing protein motifs containing acetyllysine. Although originally characterized as histone-binding proteins, it has since become clear that these domains interact with other acetylated proteins, perhaps most prominently transcription factors. The likely transient nature and low stoichiometry of such modifications, however, has made it challenging to fully define the interactome of any given bromodomain. To begin to address this knowledge gap in an unbiased manner, we carried out mRNA display screens against a bromodomain – the *N*-terminal bromodomain of BRD3 – using peptide libraries that contained either one or two acetyllysine residues. We discovered peptides with very strong consensus sequences and with affinities that are significantly higher than typical bromodomain-peptide interactions. X-ray crystal structures also revealed modes of binding that have not been seen with natural ligands. Intriguingly, however, our selected sequences are not found in the human proteome, perhaps suggesting that strong binders to bromodomains might have been selected against.

## INTRODUCTION

Biology is built on networks of protein-protein interactions that have affinities that span over 10 orders of magnitude (1). It has often been mooted that protein-protein interactions – particularly those involved in signalling networks – are not necessarily optimized for high affinity because lower affinities might provide higher sensitivity in response to environmental stimuli. For example, although SH3 domains often bind target polyproline motifs with affinities of 10–100 μM (2), K_D_s as low as 10 nM have been reported (3) demonstrating that the SH3-polyproline system is capable of affinities ∼1000-fold higher than the typical range.

Relatively weak interactions are abundant in transcriptional regulation, including in the recognition of histone post-translational modifications by ‘reader’ domains. A prominent example is the interaction between bromodomains (BDs) and linear motifs containing acetyllysine (AcK) residues. BDs are small alpha-helical bundles found in ∼40 human proteins that mainly regulate gene expression (4). These domains harbour a deep pocket that specifically recognizes AcK-containing peptide motifs in both histones and other gene regulatory proteins such as transcription factors (9-12) The affinities of these interactions lie in the range 10–200 μM (*e.g.*, (5,6))), although we have recently shown that non-native cyclic peptides containing AcK residues can achieve nanomolar and even picomolar affinities (14), highlighting that higher affinities are in principle possible.

Because of the complex and networked nature of histone-protein interactions, it is frequently difficult to unambiguously establish the cognate target(s) of a particular BD. The large number of possible non-histone acetylation events – particularly among transcription factors – and their likely low stoichiometries *in vivo* has compounded this issue, hindering the comprehensive identification of the BD interactome (13).

To being to obtain an unbiased and broad-based view of BD target sequences, we used Random non-standard Peptide Integrated Discovery (RaPID) mRNA display (15). We created libraries of 9- or 12-residue peptides containing one or two AcK residues, respectively, and screened these libraries against a well-studied BD: the *N*-terminal bromodomain (BD1) of BRD3, a member of the bromodomain and extra-terminal domain (BET) sub-family of BD proteins (**Fig. S1A**). These proteins each possess two BDs that can recognize motifs bearing either a single AcK residue or two AcK residues (di-AcK); di-AcK motifs generally have the consensus AcK-XX-AcK, where X is any amino acid (7,8)) (**Fig. S1B**).

From our screen, peptides with very strong consensus sequences were isolated from the screens and biophysical analysis of the most enriched peptide from the di-AcK screen revealed affinities for BET-family BDs of ∼1–2 μM, ∼5× tighter than the strongest known single BD-substrate interaction (16). X-ray crystal structures of the mono-AcK and di-AcK peptides bound to BRD3-BD1 revealed the structural basis for these interactions, showing that they are unlike any interactions with known physiological ligands. Unexpectedly, these consensus sequences do not appear in the human proteome, suggesting that the tightest-binding target sequences for this domain have perhaps not been selected for during evolution. This finding is in line with the idea that the ability for interactions to be rapidly regulated is an important aspect of cellular processes such as gene regulation.

## RESULTS

### RaPID screens against BRD3-BD1 select highly enriched AcK-containing sequences

To assess the substrate preferences for a BD in a comprehensive manner, we designed two DNA-encoded linear peptide libraries that used genetic code reprogramming to incorporate AcK at AUG codons (17) (**Fig. 1A** and **Fig. S2**). One library encoded peptides of the type ^*N*^AcA-X_4_-AcK-X_4_ (the 1AcK library), where X is any natural amino acid other than Trp and Met, and ^*N*^AcA is an *N*-terminally acetylated Ala. Given the ability of BET BDs to recognize sequences containing two AcK residues (most often separated by two residues), we also created a library with the composition ^*N*^AcA-X_4_-AcK-X_2_-AcK-X_4_ (the 2AcK library). We chose the first BD of the human BET-family protein BRD3 (BRD3-BD1) as a target and separately panned the two libraries against biotinylated BRD3-BD1 immobilized on streptavidin beads.

**Figure 1.**
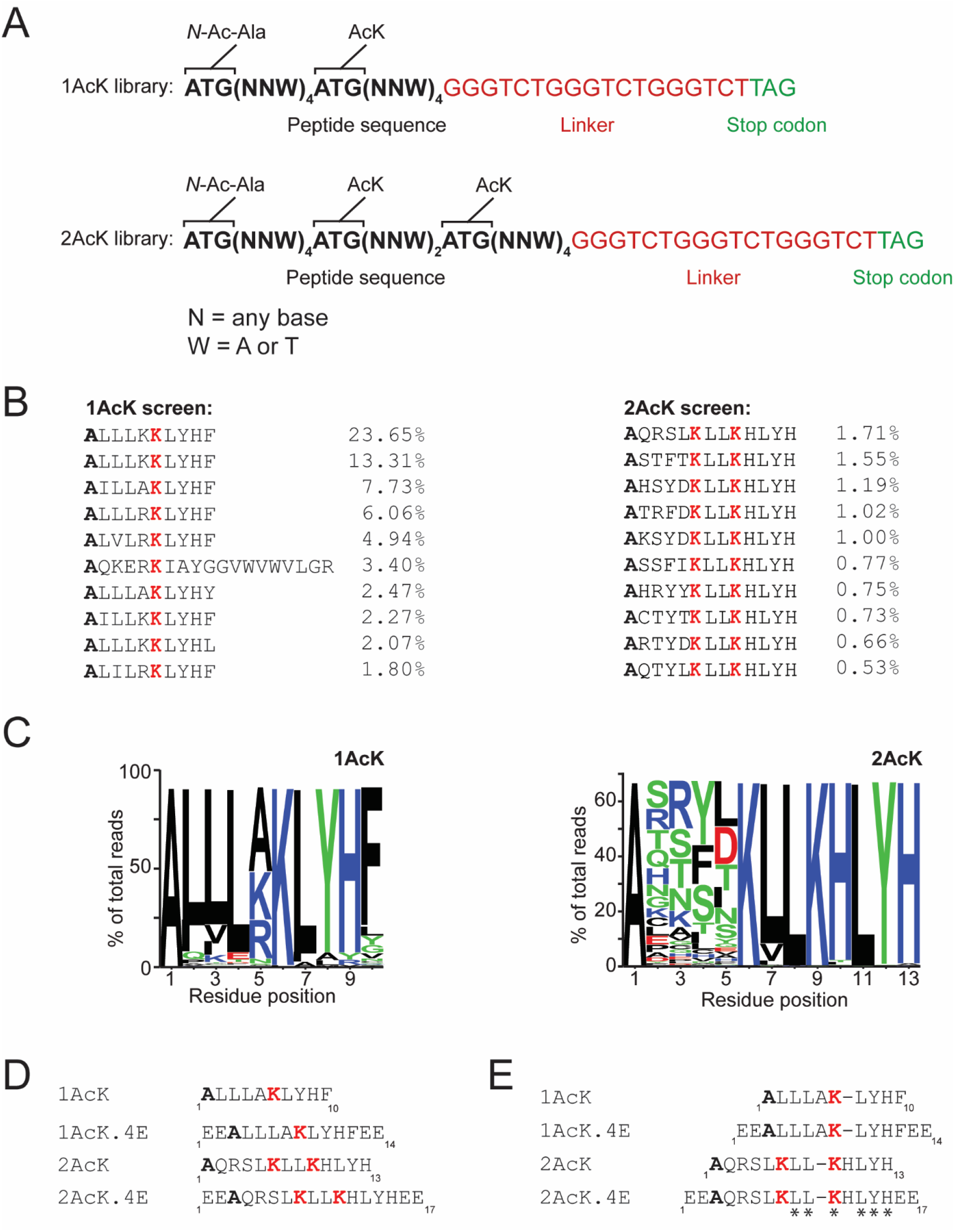
RaPID screens for BRD3-BD1 binding peptides. **A**. Library design for the 1AcK and 2AcK libraries. NNW codons encode any amino acid except Trp and Met. **B**. Alignment of the ten most highly represented sequences from each screen. The percentage of reads of each sequence is shown as a fraction of the total number of mapped reads in each screen. The top sequence from each screen was synthesized for further analysis. **C**. Sequence logo (generated from the 1000 most abundant sequences using WebLogo) showing the abundance of each amino acid in each position of the selected sequences. **D**. Peptides used for binding analysis in this study. **E**. Alignment of selected sequences used in this study. *Red* indicates acetyllysine, whereas *bold* indicates the *N*-terminal residue in the parent peptide sequence that carries an acetyl group. Positions that are conserved across the sequences used are indicated with asterisks (*) beneath the sequences.

Following four rounds of mRNA display selection, the DNA libraries generated from each round were sequenced. Clear enrichment was observed even after only two rounds, and these preferred sequences were further enriched after four rounds (**Table S1**). **Fig. 1B** shows the ten most highly represented sequences from each selection. In line with the library design, each peptide contained either one or two AcK residues. Weighted consensus sequences derived from the top 1000 sequences (which comprised >80% and >60% of total DNA sequencing reads from the 1AcK and 2AcK selections, respectively), show the emergence of strong consensus sequences in the majority of the randomized positions (**Fig. 1C**). In addition, the most enriched sequence from each selection also conforms to this 1000-sequence consensus.

Substantial similarity was also observed between the sequences selected in the two experiments (**Fig. 1D** and **1E**). Leu is highly enriched in the *N*-terminal part of each sequence and a Leu-Tyr-His sequence in the C-terminal part (**Fig. 1E**). Interestingly, although there appeared to be amino acid preferences in all positions in the single AcK library, almost no preference was observed in the positions *N*-terminal to the first AcK in the 2AcK selection (**Fig. 1B** and **1C**).

### The most enriched 2AcK sequence binds BD1 domains with high affinity

We synthesized the most enriched peptide from each selection (1AcK and 2AcK; **Fig. 1B** and **1D**) and used surface plasmon resonance (SPR) to assess their ability to bind each of the two BDs from the BET paralogues BRD2, BRD3 and BRD4, which were each immobilised on a streptavidin-coated SPR chip. The peptide 1AcK was not sufficiently soluble to allow measurements to be made; 2AcK was slightly more soluble and displayed robust and reproducible binding to each of the three BD1 domains, with affinities of ∼1–2 μM to BRD3-BD1, BRD4-BD1 and BRD2-BD1 (**Fig. 2A, Table 1**). Affinities for the BD2 domains were substantially lower with no measurable binding to BRD2-BD2 and BRD3-BD2, and a small degree of binding observed to BRD4-BD2, though an affinity could not be reliably determined.

**Table 1.** Dissociation constants for the cyclic peptides selected for study from RaPID selections against BRD2-BD2, BRD3-BD2, and BRD4-BD2.

| Peptide | $K_D$ ($\mu\text{M}$) | | | | | |
| --- | --- | --- | --- | --- | --- | --- |
|  | BRD2-BD1 | BRD3-BD1 | BRD4-BD1 | BRD2-BD2 | BRD3-BD2 | BRD4-BD2 |
| 1AcK | nd<sup>b</sup> | nd<sup>b</sup> | nd<sup>b</sup> | nd<sup>b</sup> | nd<sup>b</sup> | nd<sup>b</sup> |
| 2AcK | 1.6 ± 0.67 | 1.5 ± 0.98 | 1.2 ± 0.02 | nb<sup>c</sup> | nb<sup>c</sup> | >10 |
| 1AcK.4E | >200 | >200 | >200 | >200 | >200 | >200 |
| 2AcK.4E | 69 ± 18 | 22 ± 3.5 | 53 ± 8.7 | >200 | >200 | >200 |

<sup>a</sup>The $K_D$ values are given as the geometric mean of a minimum of three independent measurements. The standard deviation is provided.
<sup>b</sup>Not determined due to poor solubility of the peptide.
<sup>c</sup>No binding observed for the interaction.

**Figure 2.**
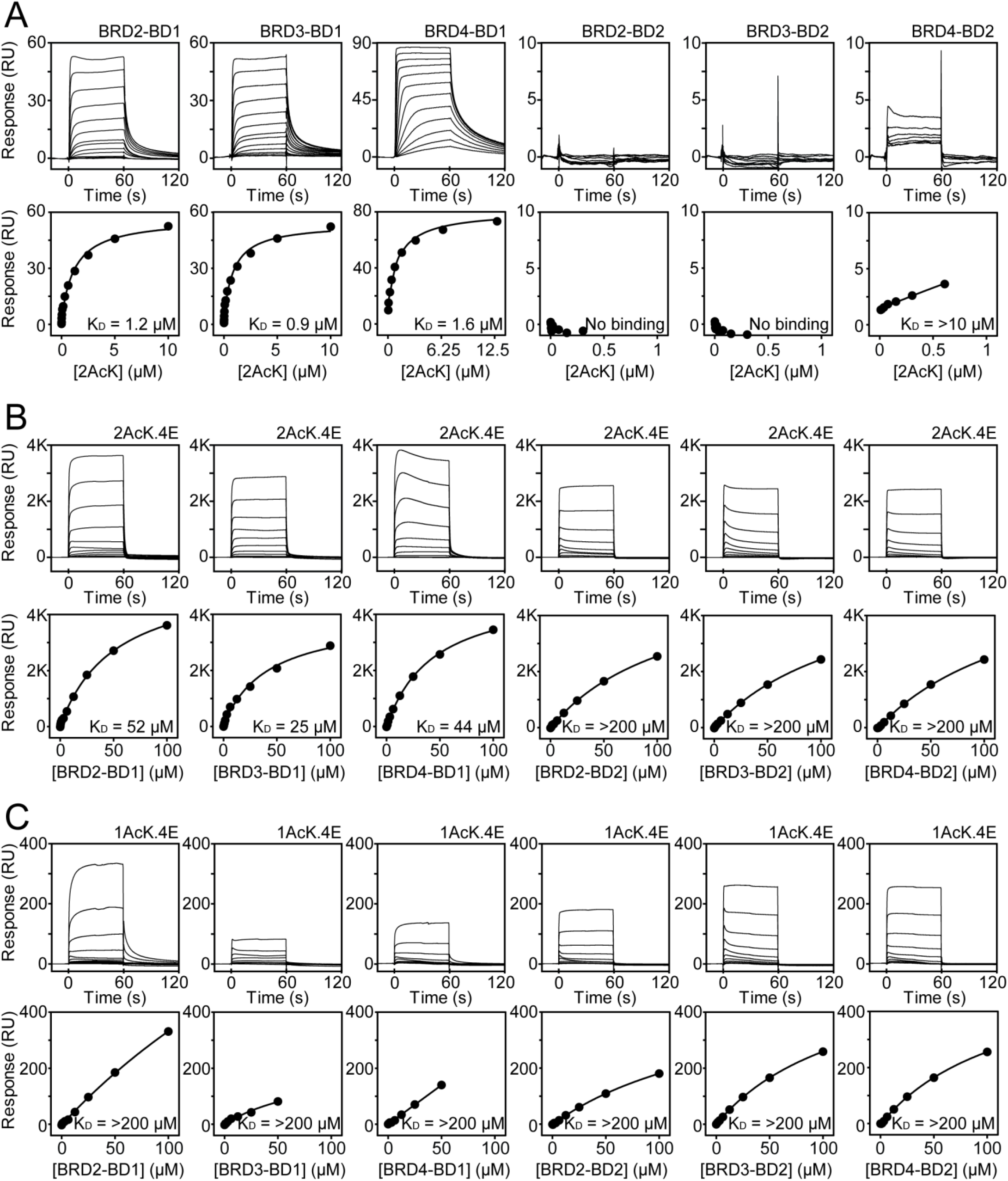
SPR analysis of BD-peptide interactions. *Top*. Typical sensorgrams are shown for one titration of each of the six indicated BDs with (**A**) 2AcK, (**B**) 2AcK.4E and (**C**) 1AcK.4E. *Bottom*. Fits to a 1:1 binding model are shown; K_D_s are indicated on each plot. Each experiment was conducted at least three times and geometric mean K_D_ values and associated uncertainties are given in Table 1.

Due to the low solubility of the original peptides, we synthesized analogues that incorporated two glutamates at both the *N*- and *C*-termini of each peptide to facilitate characterisation of the peptide-BD interactions (**Fig. 1D**). The solubility of the new peptides (1AcK.4E and 2AcK.4E) was much improved. The affinities of biotinylated variants of the glutamate-containing peptides for each BET BD were determined by immobilising the peptides to a streptavidin-coated SPR chip and titrating with the BDs. The 2AcK.4E peptide bound BD1 domains with affinities of ∼20–70 μM and BD2 domains with affinities weaker than 200 μM (**Fig. 2B, Table 1**). The glutamates thus improved solubility at the expense of affinity. Nevertheless, we could now measure binding of 1AcK.4E, which bound BD1 domains with affinities weaker than the corresponding interactions with 2AcK.4E. As none of the titrations against 1AcK.4E went to completion in our experiments, all affinities were estimated to be >200 μM (**Fig. 2C**).

### Peptides bind in the canonical AcK-binding pocket

To investigate the binding mode of these peptides, we carried out ^15^N-HSQC chemical shift perturbation experiments using both 1AcK.4E and 2AcK.4E. Furthermore, to explore whether the addition of solubilising glutamates had altered the peptide-BD binding mode, we also tested 2AcK, which was sufficiently soluble to perform these experiments using low concentrations of BDs (e.g., by using ∼30 μM ^15^*N*-labelled BD, the addition of eight molar equivalents of peptide could be achieved). ^15^N-HSQC spectra were recorded following each addition of peptide into uniformly ^15^N-labelled BRD3-BD1 or BRD3-BD2.

In the titration of 2AcK into BRD3-BD1, widespread but selective reductions in signal intensity were observed (**Fig. 3A**). The pattern of changes indicates that the interaction is in slow-to-intermediate exchange on the chemical shift timescale, consistent with the affinity observed by SPR. In contrast, and in line with the lower affinities measured by SPR, titrations with 2AcK.4E were in the intermediate-fast exchange regime, displaying perturbations to both signal intensity and position (**Fig. S3A**). To map the binding site of both peptides, we quantified the percentage decrease in intensity of each signal before and after the titration and plotted these changes against residue number (**Fig. 3B** and **Fig. S3B**). Most changes were common to both peptides and mapping of these changes onto the structure of BRD3-BD1 indicated a common BD-binding surface – one that directly overlaps the canonical AcK-binding pocket. Thus, both peptides bind to the native AcK binding site and, although the introduction of flanking glutamates in 2AcK.4E reduces the affinity of the peptide, it does not alter its binding site.

**Figure 3.**
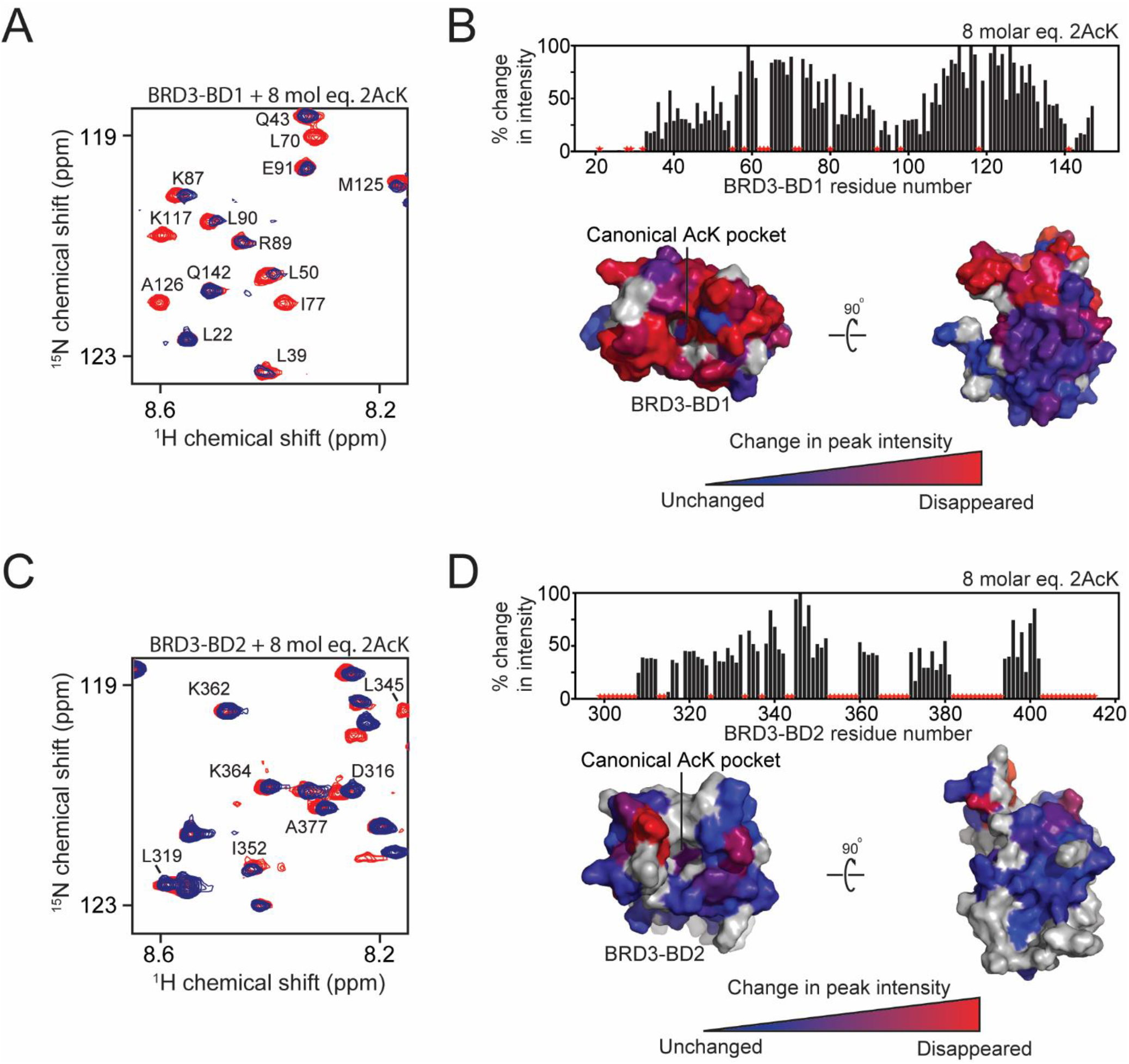
^15^N-HSQC titrations for 2AcK into the BDs of BRD3 and structural mapping of the interactions. For all ^15^N-HSQC titrations, BDs were used at a concentration of ∼30 µM and the assignments and direction of movement of some signals are indicated. Residues that could not be assigned in the HSQCs are indicated by red stars in the graphs plotting intensity change and in grey on the structure of BDs. **A**. ^15^N-HSQC spectra of BRD3-BD1 alone (*red*) and in the presence of 8 molar equivalents of 2AcK (*blue*). **B**. *Top*: Quantitation of change in ^15^N-HSQC signal intensity for each signal following the addition of the indicated amount of 2AcK to BRD3-BD1. *Bottom:* Intensity changes from the graph above mapped onto the structure of BRD3-BD1. The magnitude of the change is represented by a *blue–red* colour gradient (red indicates the largest change). **C**. ^15^N-HSQC spectra of BRD3-BD2 alone (*red*) and in the presence of 8 molar equivalents of 2AcK (*blue*). **D**. Top: Quantitation of change in ^15^N-HSQC signal intensity for each signal following the addition of the indicated amount of 2AcK to BRD3-BD2. *Bottom*: Intensity changes from the graph above mapped onto the structure of BRD3-BD2. The magnitude of the change is represented by a *blue–red* colour gradient (*red* indicates the largest change).

In the titrations of 2AcK and 2AcK.4E against BRD3-BD2, substantially smaller chemical shift perturbations were observed, indicating both interactions are in fast exchange on the chemical shift timescale and suggestive of a substantially weaker binding affinity for BRD3-BD2 than BRD3-BD1 (**Fig. 3C** and **Fig. S3C**). We quantified these changes in peak intensity and mapped them onto the BRD3-BD2 structure. As for BRD3-BD1, these changes mapped to the canonical AcK binding pocket on the BD2 (**Fig. 3D** and **Fig. S3D**).

The ^15^N-HSQC titrations of 1AcK.4E against BRD3-BD1 and BRD3-BD2 revealed chemical shift perturbations to largely the same peaks observed in the corresponding titrations with 2AcK and 2AcK.4E (**Fig. S4A** and **Fig. S4C**). However, fewer peaks changed and the observed changes were smaller, in line with the weak (>200 µM) affinities measured for these interactions by SPR. Mapping the intensity changes onto structures of BRD3-BD1 and BRD3-BD2 once again revealed binding largely to the same site on the BDs as for the 2AcK and 2AcK.4E (**Fig. S4B** and **Fig. S4D**).

Since we observed more substantial peak perturbations in our titrations using 2AcK and 2AcK.4E compared to 1AcK, we went on to explore their interactions with the BDs of BRD2 and BRD4. When compared to BRD3-BD1, similar sets of peaks in BRD2-BD1 and BRD4-BD1 were perturbed on addition of both peptides, though the overall magnitude of these changes was smaller (Fig. 4A-D and **Fig. S5A-D**). As observed with BRD3-BD1, the 2AcK peptide induced chemical shift perturbations to the BD1s that were reminiscent of a slow-intermediate exchange regime whereas 2AcK.4E displayed an intermediate-fast exchange regime with both signal intensity changes and movement observed. Of the peptide titrations against BD2 domains, the largest peak intensity and chemical shift perturbations were observed with 2AcK titrated against BRD4-BD2 (Fig. 4G and Fig. 4H), whereas titrations with 2AcK.4E resulted in comparable changes across all BD2 domains (**Fig. S3C-D** and **Fig. S5E-F**) These results are in line with the SPR-derived affinities. In all cases we mapped the chemical shift perturbations onto the structures of the BDs, revealing that the binding site was common to all BDs tested (Fig 4 and Fig S5).

**Figure 4.**
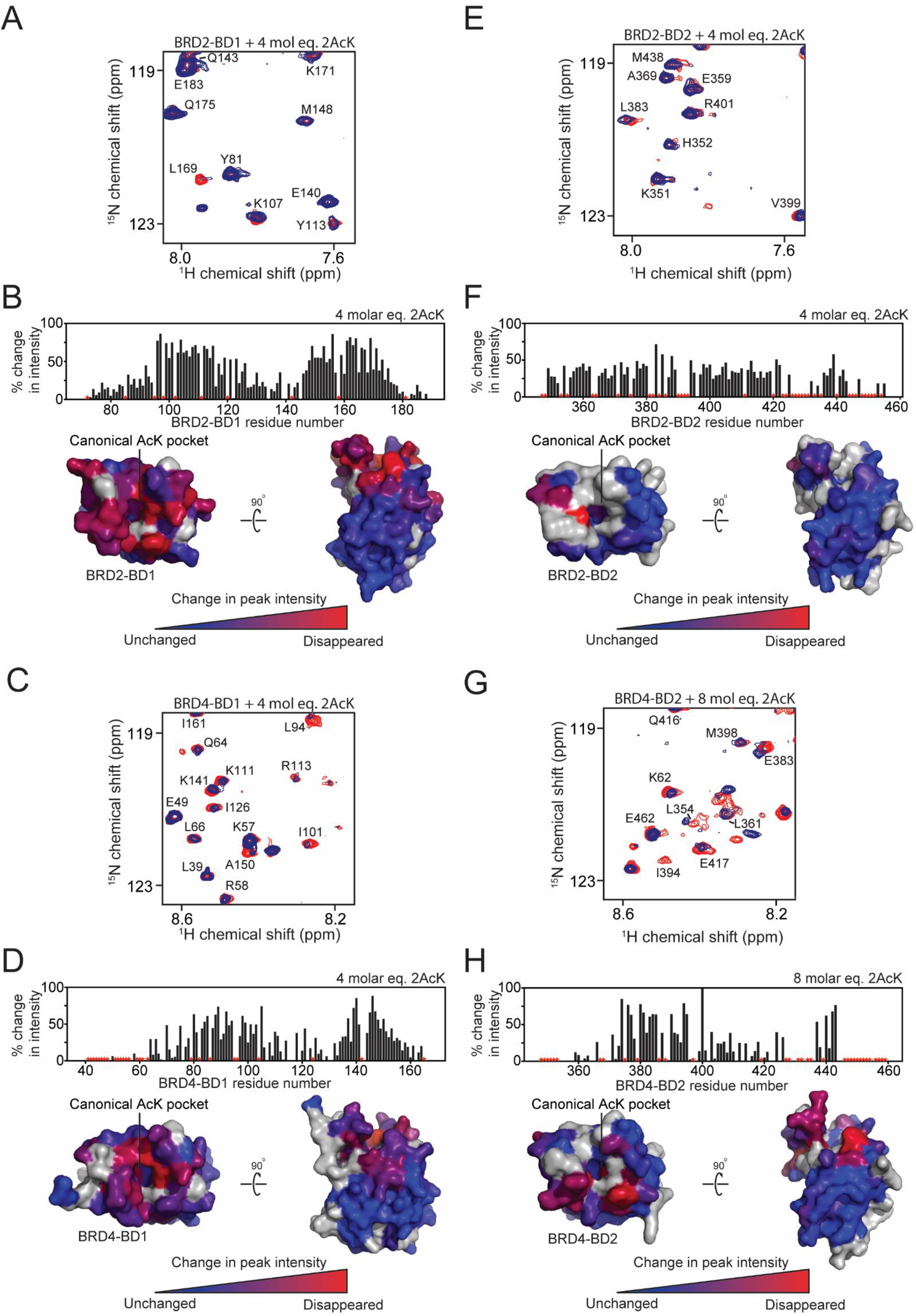
Peptide 2AcK binds the BDs from BRD2 and BRD4 at the canonical binding site. For all ^15^N-HSQC titrations, BDs were used at a concentration of ∼30 µM and the assignments and direction of movement of some signals are indicated. Residues that could not be assigned in the HSQCs are indicated by red stars in the graphs plotting intensity change and in grey on the structure of BDs. **A**. ^15^N-HSQC spectra of BRD2-BD1 alone (*red*) and in the presence of 4 molar equivalents of 2AcK (*blue*). **B**. *Top*: Quantitation of change in ^15^N-HSQC signal intensity for each signal following the addition of the indicated amount of 2AcK to BRD2-BD1. *Bottom*: Intensity changes from the graph above mapped onto the structure of BRD2-BD1. The magnitude of the change is represented by a *blue–red* colour gradient (*red* indicates the largest change). **C**. ^15^N-HSQC spectra of BRD4-BD1 alone (*red*) and in the presence of 4 molar equivalents of 2AcK (*blue*). **D**. *Top*: Quantitation of change in ^15^N-HSQC signal intensity for each signal following the addition of the indicated amount of 2AcK to BRD4-BD1. *Bottom*: Intensity changes from the graph above mapped onto the structure of BRD4-BD1. The magnitude of the change is represented by a *blue–red* colour gradient (*red* indicates the largest change). **E**. ^15^N-HSQC spectra of BRD2-BD2 alone (*red*) and in the presence of 4 molar equivalents of 2AcK (*blue*). **F**. *Top*: Quantitation of change in ^15^N-HSQC signal intensity for each signal following the addition of the indicated amount of 2AcK to BRD2-BD2. *Bottom*: Intensity changes from the graph above mapped onto the structure of BRD2-BD2. The magnitude of the change is represented by a *blue–red* colour gradient (*red* indicates the largest change). **G**. ^15^N-HSQC spectra of BRD4-BD2 alone (*red*) and in the presence of 8 molar equivalents of 2AcK (*blue*). **H**. *Top*: Quantitation of change in ^15^N-HSQC signal intensity for each signal following the addition of the indicated amount of 2AcK to BRD4-BD2. *Bottom*: Intensity changes from the graph above mapped onto the structure of BRD4-BD2. The magnitude of the change is represented by a *blue–red* colour gradient (*red* indicates the largest change).

Taken together our NMR titration data indicate that all three peptides bind all tested BDs in the canonical AcK binding pocket, but with a range of affinities, and that the addition of solubilising glutamates does not substantially alter this binding site.

### 2AcK binds BRD3-BD1 via an unprecedented mechanism

We next determined X-ray crystal structures for complexes formed between BRD3-BD1 and both 2AcK (1.5 Å resolution, PDB ID: 7TO8, **Table S2**) and 2AcK.4E (1.6 Å resolution, PDB ID: 7TO9, **Table S2**). In line with our NMR data, the two structures are essentially identical and here we will describe only the former. As shown in Fig. 5, two copies of BRD3-BD1 bind a single peptide. The two BDs take up an essentially identical conformation to each other and to other reported structures of the domain. The peptide forms two regular turns of α-helix, a conformation that has not been observed in any known natural BD ligand (from a total of >30 structures). The two BDs are related by a 2-fold non-crystallographic axis that runs perpendicular to the long axis of the peptide α-helix (**Fig. S6**). This rotation maps the helix directly onto itself, except in the opposite direction. Both orientations of the peptide were found equally in the crystal structure. The symmetry orientation also caused the two pocket-binding residues to swap their positions.

**Figure 5.**
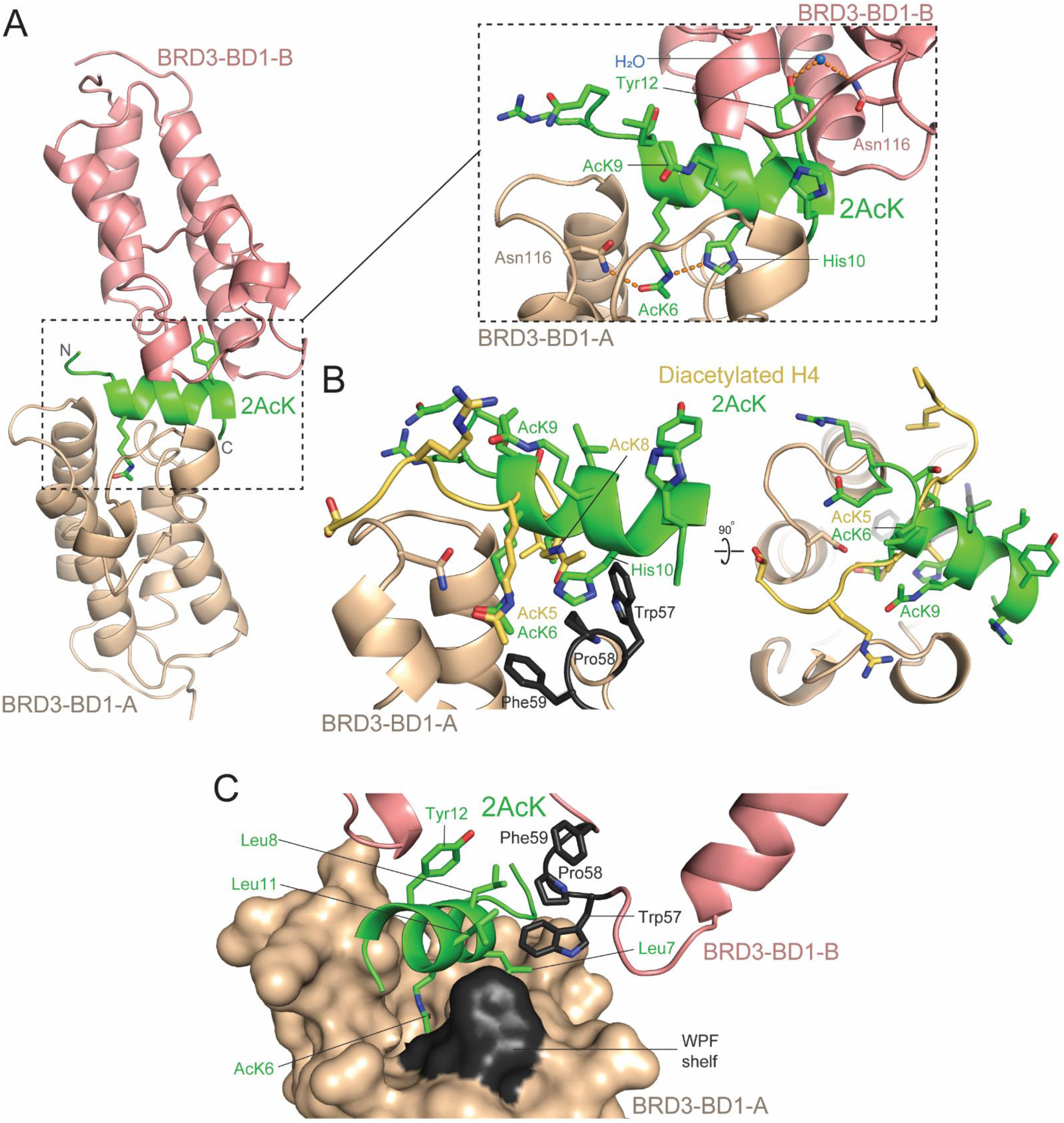
Structural basis for the BRD3-BD1:2AcK interaction. **A**.Structure of the BRD3-BD1:2AcK complex (PDB ID: 7TO8). The two molecules of BRD3-BD1, are shown in *wheat* (A) and *salmon* (B), and the peptide is shown in *green*. A close-up of the interaction is shown in the *dashed square panel*. The two AcK residues are labelled, including AcK6 that forms a hydrogen bond with Asn116 of BRD3-BD1-A, as is the Tyr12 residue that occupies the binding pocket and forms a water mediated hydrogen bond with Asn116 of BRD3-BD1-B. Hydrogen bonds are indicated by the *orange dashed lines*. **B**. Comparison of the binding of 2AcK to BRD3-BD1 (*green*) with a diacetylated histone H4 peptide (*yellow*) bound to BRD3-BD1 and BRD4-BD1, respectively. Peptide position based on superposition of BRDs molecules (PDB ID: 3UVW) (5).A view of the AcK insertion angle (*left*) and backbone conformation (*right*). **C**. Interactions of leucine residues in 2AcK with the WPF shelf (*black*) of both BRD3-BD1 molecules in complex with the peptide.

In the structure, one BD binds the *N*-terminal AcK (AcK6) of the peptide in the canonical pocket, whereas the corresponding pocket in the second BD is occupied by Tyr12 (Fig. 5A, *inset*, numbering is as in Fig. 1D). The acetyl group of AcK6 forms the same hydrogen bond with the sidechain of Asn116 that is common to all characterized BD-peptide interactions. Likewise, the overall position of AcK6 in the pocket is the same as that observed for complexes with native ligands such as AcK5 of histone H4 (Fig. 5B). In the second BD, hydroxyl group of Tyr12 forms water mediated hydrogen bond also to Asn116, (Fig. 5A, *inset*). The total buried surface area at BD-peptide interface involving AcK6 is 1320 Å^2^ and 1010 Å^2^ (for the interaction involving Tyr12), which is significantly larger than area buried in, for example, the complex formed between BRD4-BD1 and the diacetylated (on K5 and K8) histone H4 ligand (690 Å^2^; PBD ID: 3UVW)(5). In contrast to large surface contact between the peptide and each BD, the two BD chains make very few inter-domain contacts, burying a total of only 260 Å^2^ of surface area.

In structures of BET-family BDs bound to diacetylated peptides from the native targets histone H4, as well as the transcription factors GATA1, E2F1 and Twist (7,9),11)), the peptide backbone is extended and runs at roughly 90° to the long axis of the 2AcK helix (Fig. 5B). This difference allows 2AcK to place a histidine (His10) into a portion of the binding pocket that is not utilized by native partners and furthermore to form a hydrogen bond with the Nε of the amide group of AcK6 (Fig. 5A, *inset*). An arrangement of this type is not possible with native ligands because of the position of the ligand backbone (Fig. 5B).

In complexes with diacetylated native ligands, the more C-terminal of the two AcK residues forms van der Waals interactions with the so-called ‘WPF’ shelf on the edge of the binding pocket (Fig. 5B and **Fig. S1B**). In the 2AcK complex, however, the corresponding AcK (AcK9) occupies a very different position (due to the peptide helical conformation) and is not in contact with the WPF shelf of either of the two BDs (Fig. 5B). Instead, Leu7 and Leu11 forms contact with the WPF shelf of the AcK6-binding BD (Fig. 5C and **Fig. S6B**). These residues, together with Leu8, simultaneously interact with the WPF shelf of the second BD. It is worth mentioning that positions of Leu7 and Leu11 are swapped due to the rotational symmetry of the complex.

Finally, we note that sidechains of residues *N*-terminal to AcK6 were either not in contact with the BD or did not display any electron density, suggesting that they were likely not well ordered. This observation is consistent with the lack of sequence preferences for these *N*-terminal residues.

### The 1AcK.4E binding mode partially mimics 2AcK and previously discovered unnatural BD peptide ligands

We determined the X-ray crystal structure of BRD3-BD1 bound to 1AcK.4E (1.9 Å resolution, PDB ID: 7TO7, **Table S2**). The overall architecture of the complex closely resembles the BRD3-BD1:2AcK structure with two BDs binding a helical peptide and all three chains in the same relative spatial arrangement (Fig. 6A). The two copies of the BD are again related by a 2-fold non crystallography axis observed for the 2AcK structures. Again the electron density for the peptide is best modelled at 50% occupancy in the two symmetrical orientations (**Fig. S7A**). Furthermore, the binding pocket of one BD accommodates an AcK8 residue, while the other binds a Tyr10 (Fig. 1D, Fig. 6A *inset*, **Fig. S7B**). Compared to the 2AcK structures, slightly less surface area is buried while forming the complex (1100 and 1000 Å^2^, for the AcK-mediated and Tyr-mediated interactions, respectively, and 260 Å^2^ between the two BDs). Although the 1AcK.4E sequence differs significantly and sequential distance of the two pockets binding residues is smaller, compared to 2ACK, the overall peptide placing remains the same. Non ideal distance of the two main binding residues, smaller by one α-helix turn, resulted in different conformation of Ack8 in the binding pocket (Fig. 6C). We have observed this “diagonal” conformation in binding pocket for helical cyclic peptides (such as 4.2A) discovered in RaPID screens against BET bromodomains, previously (14) (Fig. 6B). In parallel with those structures, AcK8 of 1AcK.4E forms a water-mediated hydrogen bond with Asn116 (as does Tyr10; Fig. 6A *inset*).

**Figure 6.**
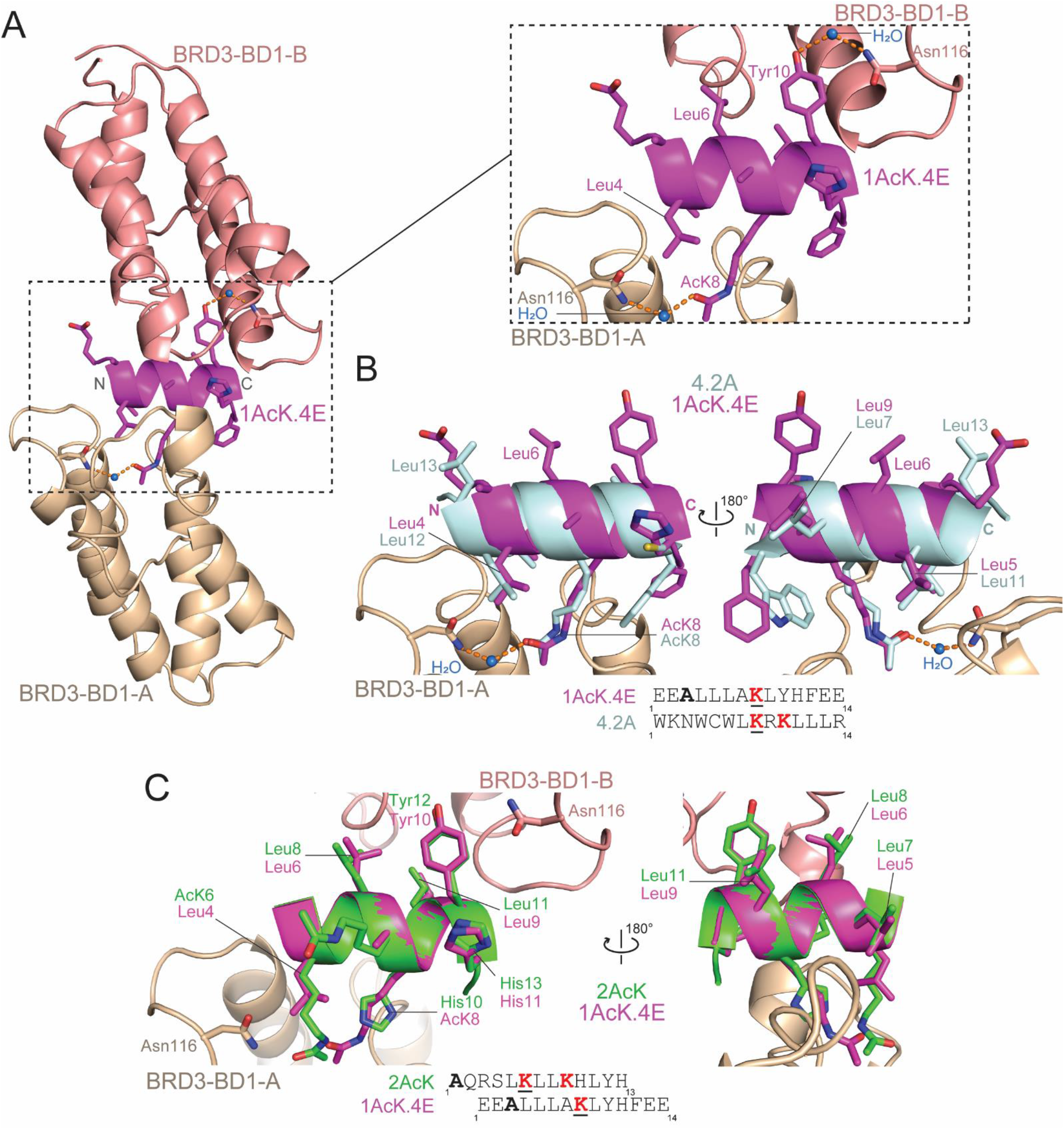
Structural basis for the BRD3-BD1:1AcK.4E interaction. **A.** Structure of the BRD3-BD1:1AcK.4E complex (PDB ID: 7TO7). The two copies of BRD3-BD1 are shown in wheat (A) and salmon (B), and the peptide is shown in magenta. A close-up of the interaction is shown in the dashed square panel. AcK8 and Tyr10, which each occupy the binding pocket of one of the BDs, are labelled, as is the water molecule (*blue* sphere) that forms hydrogen bonds bridging Asn116 and the pocket binding residue (either AcK8 or Tyr10). The hydrogen bonds are indicated by the orange dashed lines. B. Structure of the peptide 4.2A (pale cyan; crystallised in complex with BRD4-BD1, PDB ID: 6ULV) overlayed onto the structure of BRD3-BD1-A:1AcK.4E (14). The angle of entry of AcK8 in 4.2A into the binding pocket mirrors that of AcK8 in 1AcK.4E. The overlay shows the conserved locations of leucine residues that contact the WPF shelf (**Fig. S7B**). Leu7 and Leu11 of 4.2A map onto Leu9 and Leu5 of 2AcK, respectively. C. Overlay of 1AcK.4E (magenta) with 2AcK.1 (green). The backbone conformations and positions and identities of many sidechains are closely matched despite the differing positions of the pocket-binding AcK residues. The underlined red AcK residues are the ones that engage the canonical binding pocket.

Comparison of the 1AcK.4E and 2AcK structures reveals that five residues make essentially identical interactions in the two structures (Fig. 6C): the pocket-binding tyrosine (Tyr10 in 1AcK.4E; Tyr12 in 2AcK), the three leucines that pack against the WPF shelf (Leu5, Leu6 and Leu9 in 1AcK.4E; Leu7, Leu8 and Leu11 in 2AcK), and the histidine that immediately follows the tyrosine (His11 in 1AcK.4E; His13 in 2AcK). Strikingly, 4.2A also displays two of the three leucines in the same positions (Leu7 and Leu11), with the aliphatic portion of an AcK sidechain occupying the third position to pack against the WPF shelf (Fig. 6B). It is also notable that this similarity in positioning of these residues occurs despite the fact that the 4.2A helix runs in the opposite direction to the peptides discovered in this study (Fig. 6B).

Finally, we assessed the conformational preferences of 2AcK.4E by NMR spectroscopy. Proton chemical shifts are most consistent with the peptide being intrinsically disordered and two-dimensional NOESY and ROESY data display very few of the NOE patterns (e.g., an extended series of HN-HN NOEs between sequential residues) that are diagnostic of helical structure (**Fig. S8**). It is therefore likely that the helical structure observed in the complexes arises from conformational selection from a rarely populated conformer.

## DISCUSSION

Reported natural ligands of the BET bromodomains have diverse sequences and affinities ranging from the tens to hundreds of micromolar. In contrast, we recently derived a set of cyclic peptide ligands that bind to BET BDs with binding affinities in the nanomolar and high picomolar range (14). In the current study, we therefore sought to determine whether linear sequences could also achieve higher affinities than known biological ligands and therefore might represent an as yet unrecognized class of natural BET BD targets.

In this work, we devised and tested a strategy that we hypothesized could be used to deliver a more comprehensive view of preferred BD binding motifs. We used RaPID mRNA display to screen two peptide libraries, each comprising ∼10^12^ peptides and harbouring either one or two AcK residues. The AcKs were separated by two residues to mimic the arrangement found most frequently in natural diacetylated BET ligands. Although we envisaged that our screens might provide a broad overview of the substrate preferences of BRD3-BD1, each screen converged rapidly on one major consensus sequence – and these sequences were similar though not identical between the two screens. Despite what appeared to be an unusually strong preference for these sequences, their affinities for BET bromodomains (as high as ∼1 μM for 2AcK) were still several orders of magnitude weaker than those observed for our previously identified cyclic peptides. The affinity of 2AcK for BRD3-BD1 was nonetheless higher than all other measured natural ligands.

Unexpectedly, however, a search for these sequences in the human proteome, or even for the ‘core’ sequences KLLKHLYH (from 2AcK) and LLLXKLYHF (from 1AcK, where X can be any amino acid), revealed no exact matches. Although this observation is not definitive evidence that the sequence has been selected against during evolution, it is nonetheless surprising given the large number of proteins to which the BET bromodomains have been shown to bind.

### A new BET BD binding mode for linear peptides

The binding modes of our selected peptides also differed significantly from previously characterised natural ligand-BET BD binding interactions. A structure of 2AcK bound to BRD3-BD1 revealed that, although the more *N*-terminal AcK binds with a conformation closely resembling the corresponding residue in native substrates, the second AcK contacts a completely different surface (Fig. 5). The size of the interface made with each BD is also substantially greater than that observed for native partners with as many as nine residues from the peptide contacting both BDs in the crystal structure.

Both screens yielded peptides that formed a regular α-helix. This conformation is not adopted by any known naturally occurring BET target sequences, although several of our previously isolated cyclic peptide ligands have helical structures (14). In all cyclic peptides that we structurally characterized, the helix backbone overlays closely with the helix observed in this current study and can run in either direction. Strikingly, bidirectionality of binding to BDs within a single peptide, as observed in this study, is new. Furthermore, the pocket-binding AcK in all previously discovered helical peptides had the ‘diagonal’ entry angle observed for 1AcK.4E in the current study – in contrast to the perpendicular angle observed for the 2AcK peptide and for our tightest-binding cyclic peptides.

The affinity of 2AcK for BET-BD1 domains is lower than the affinities observed for our helical cyclic peptides by a factor of ∼50, broadly suggesting that macrocycle formation can boost the achievable affinity for this interaction modality by one to two orders of magnitude, most likely due to the reduction in conformation entropy that needs to be lost upon complex formation. This idea is consistent with our observation that 2AcK.4E is largely disordered in solution (14).

### Screening with a monoacetylated library suggests that positioning of the AcK in the helix is important for high-affinity binding

The library design for our 1AcK screen placed the ‘warhead’ AcK near the centre of the peptide – residue six in a ten-residue sequence. This arrangement precluded the selection of peptides with the same architecture as 2AcK, because all seven residues C-terminal to the pocket-binding AcK in 2AcK make direct contact with the BDs, and only four residues are available in the 1AcK peptide. Instead, the most highly selected sequences from the 1AcK screen retain five of the key binding residues (Leu7, Leu8, Leu11, Tyr12 and His13) in the same relative positions but ‘shift’ the AcK in the C-terminal direction by one helical turn.

The resulting diagonal entry angle for this AcK leads to formation of a water-bridged hydrogen bond between the AcK and Asn116, in contrast to the direct hydrogen bond formed by 2AcK. Comparison of the affinities measured for 1AcK.4E and 2AcK.4E suggests that the difference in AcK binding mode is worth at least three-fold in affinity for binding to BD1 domains. In contrast, binding to BD2 domains is similar for 1AcK and 2AcK peptides, although the structural basis for this distinction is not clear.

Strikingly, both the 1AcK and 2AcK peptides also use a Tyr to partially occupy the binding pocket of the second BD in the complex. While there are previous instances of Tyr in the binding pocket, all examples had a ‘primary’ AcK in the pocket as well with the Tyr stabilising this AcK via a water mediated bridge (6). The direct mode of Tyr interaction with Asn116 we report here, albeit through a water mediated hydrogen bond, has not been described before. Again, it is noteworthy that this mode of interaction with Asn116 is mirrored by the AcK in the 1AcK peptide here and also the AcK in the 4.2A cyclic peptide from (14) – raising the possibility that such interactions might occur in vivo, though most likely with lower affinity.

In conclusion, given that the affinity observed for 2AcK is markedly stronger than that observed for many known natural ligands of BET proteins, our data suggest that lower affinities might be better suited to the signalling role played by these proteins. Similar arguments have been made for other protein-based signalling systems and high-throughput screening approaches such as mRNA display represent a useful tool for exploring such hypotheses.

## EXPERIMENTAL PROCEDURES

### Protein expression and purification

The bromodomains from human BRD2 (BD1: 65-194; BD2: 347-455), BRD3 (BD1: 25-147; BD2: 307-419), and BRD4 (BD1: 42-168; BD2: 348-464) were expressed in E. coli from pQE80L-Navi (Qiagen) as *N*-terminally His-tagged and biotinylated proteins for SPR. For other experiments, they were expressed from pGEX-6P (Cytiva) as *N*-terminal GST-fusion proteins. Expression and purification, via affinity chromatography and size-exclusion chromatography, of unlabelled and uniformly ^15^N-labelled proteins were carried out as described previously (14).

### RaPID screening

We synthesised two linear libraries from oligos purchases from Eurofins Genomics K.K (Japan). Each DNA sequence contained the standard RaPID T7 polymerase binding site, a ribosome binding site, a randomised peptide coding region, a (Gly-Ser)_3_ linker, an amber stop codon and a sequence for puromycin ligation. These libraries were prepared following standard protocols, transcribed to RNA and ligated with puromycin for use in the two selections.

1AcK

library:

TAATACGACTCACTATAGGGTTGAACTTTAAGTAGGAGATATATCCATG(NNW)_4_

ATG(NNW)_4_GGGTCTGGGTCTGGGTCTTAGGTAGGTAGGCGGAAA

2AcK library: TAATACGACTCACTATAGGGTTGAACTTTAAGTAGGAGATATATCCATG(NNW)_4_ATG(N NW)_2_ATG(NNW)_4_GGGTCTGGGTCTGGGTCTTAGGTAGGTAGGCGGAAA

Using these linear libraries, RaPID screens were performed as previously reported (14). The first-round translation was performed on an 80 µL scale and subsequent rounds on a 5 µL scale. For codon reprogramming, the 3,5-dinitrobenzyl ester of N^ε^-acetyl-lysine was synthesised as previously described and aminoacylated onto tRNA^Asn^_CAU_ using dFx (2 h, RT, standard aminoacylation conditions) whilst tRNA^fMet^_CAU_ was aminoacylated with N- (acetoxy)-L-alanine (Ac-L-Ala), via the 3,5-dinitrobenzyl ester using dFx (72 h, 4 °C, standard aminoacylation conditions). Flexizymes were prepared as described previously (18).

Peptide sequences from deconvoluted Illumina sequencing data are presented in **Table S1**.

### Peptide synthesis

Peptides 2AcK and 1AcK were purchased from Ontores. Peptides 2AcK.4E and 1AcK.4E were synthesised as C-terminal amides using standard fluorenylmethyloxycarbonyl (Fmoc)-strategy solid-phase chemistry using a SYRO I automated synthesiser (Biotage). Fmoc-protected amino acids (Fmoc-Xaa-OH), coupling reagents and resins were purchased from Mimotopes or Novabiochem. N,N-dimethylformamide (DMF) for peptide synthesis was purchased from RCI.

The resin (89 mg, 50 µmol, 0.56 mmol g^-1^, 1 eq.) was treated with 40 % piperidine in DMF (800 µL) for 4 min, drained, then treated with 20 vol.% piperidine in DMF (800 µL) for 4 min, drained, and washed with DMF (4 × 1.6 mL). The resin was then treated with a solution of Fmoc-Xaa-OH or chloroacetic acid (200 µmol, 4 eq.) and Oxyma (220 µmol, 4.4 eq.) in DMF (400 µL), a 1 wt.% solution of 1,3-diisopropyl-2-thiourea in DMF (400 µL), followed by a solution of DIC (200 µmol, 4 eq.) in DMF (400 µL). Coupling of Fmoc-Cys(Trt)-OH and Fmoc-His(Trt)-OH were carried out at 50 ºC for 30 min. All other coupling reactions were conducted at 75 ºC for 15 min. The resin was then drained and washed with DMF (4 × 800 µL) before being treated with a solution of 5 vol.% Ac_2_O and 10 vol.% *i*Pr_2_NEt in DMF (800 µL) for 6 min at room temperature, drained and washed with DMF (4 × 800 µL).

The resin was thoroughly washed with CH_2_Cl_2_ (4 × 7 mL) before being treated with 90:5:5 v/v/v TFA:tri*i*sopropylsilane:H_2_O and shaken at room temperature for 2 h. The resin was filtered and the filtrate concentrated under a stream of nitrogen before addition of diethyl ether (40 mL). The peptide was pelleted by centrifugation (3 min, 2 ºC, at 6500 RCF) and the ether was decanted.

Preparative reversed-phase high performance liquid chromatography (HPLC) was performed using a Waters 600E multi-solvent delivery system with a Rheodyne 7725i injection valve (5 mL loading loop) with a Waters 500 pump and a Waters 490E programmable wavelength detector operating at 214 nm and 280 nm. Preparative reversed-phase HPLC was performed using a Waters X-Bridge^®^ C18 OBD™ Prep Column (5 µm, 19 × 150 mm) at a flow rate of 15 mL min^-1^ using a mobile phase of 0.1% TFA in water (solvent A) and 0.1% TFA in MeCN (solvent B) on linear gradients, unless otherwise specified. Purity of peptides was determined by mass spectrometry recorded on a Shimadzu 2020 (ESI) mass spectrometer operating in positive and negative mode (**Fig. S9** and **S10**). Gradient grade acetonitrile (MeCN) for liquid chromatography was purchased from Merck.

### Surface plasmon resonance

Measurements were conducted on a T200 (Cytiva) and data analysed using the Biacore Insight Evaluation Software. Experiments were performed at 4°C in multi cycle kinetics mode. For experiments measuring the binding of the 2AcK peptide, biotinylated BET BDs were immobilized on a biotin CAP chip (Cytiva) with a target density of ∼1000-2000 RU and peptide was flowed over the chip. A buffer comprising 20 mM MES pH 6.5, 150 mM NaCl, 0.05% (v/v) Tween-20 was used as the running buffer. For experiments measuring the binding of 1AcK.4E and 2AcK.4E, C-terminally biotinylated variants of the peptides were immobilized on a CM5 chip (Cytiva), that had been amine-coupled with streptavidin, to a target density of ∼50-100 RU and unlabelled BET BDs were flowed over the chip. A buffer comprising 20 mM HEPES pH 7.5, 150 mM NaCl, and 0.05% (v/v) Tween-20 was used as the running buffer.

### NMR spectroscopy

NMR samples of BET BDs were prepared at ∼25-100 µM for peptide titrations. Spectra were acquired at 298 K using Bruker Avance III 600- or 800-MHz NMR spectrometers fitted with TCI probe heads and using standard pulse sequences from the Bruker library. TOPSPIN3 (Bruker) and NMRFAM-SPARKY were used for spectral analysis (19). Spectra were internally referenced to 10 μM 4,4-dimethyl-4-silapentane-1-sulfonic acid. Chemical shift perturbation (CSP) experiments were performed by collecting ^15^N-HSQC spectra of ^15^N-labelled bromodomains before and after titration of unlabelled peptide into the samples.

Chemical shift assignments were taken from previous work (14) and CSP plots were generated using the following approach. A reference HSQC was recorded without the ligand present (ref HSQC), and then the ligand was added and a second HSQC (shift HSQC) recorded. The two HSQCs were peak picked using the APES algorithm in NMRFAM-Sparky to generate peak lists. The distance between each peak on the ref HSQC and all peaks on the shift HSQC was computed as follows:

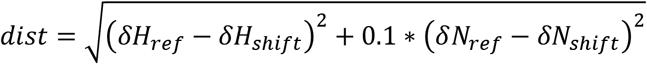

For intensity-based plots, peak height values were extracted from the ref HSQC and shift HSQC, and the values were normalized against values from peaks that were invariant throughout the titration. Percentage changes between ref HSQC and shift HSQC were then calculated and used for figure plots.

### X-ray crystallography

All crystallization studies described in this study were performed at 18 ºC using commercial 96-well crystallization screens in a sitting-drop vapor-diffusion format. Purified BRD3-BD1 (15 mg/mL) was mixed with ∼5 molar equivalents of peptide and the complexes were incubated for 0.5 h before being dispensed into MRC two-drop chamber, 96-well crystallization plates using a Mosquito crystallization robot (TTP Labtech). Complex:precipitant ratios of 1:1 and 2:1 were trialled ensuring that a final drop volume of 300 nL was maintained. Crystals of the BRD3-BD1:peptide complexes typically appeared within 1–2 days after setting up the experiments. The BRD3-BD1:2AcK complex crystallized in a solution containing 0.1 M MMT pH 6.0 and 25% (w/v) PEG1500. The BRD3-BD1:2AcK.4E complex crystallized in a solution containing 0.1 M HEPES pH 7.5, 0.2 M NaCl, and 10% (v/v) 2-propanol. The BRD3-BD1:1AcK complex crystallized in the solution containing 0.1 M PTCP pH 4.0 and 25% (w/v) PEG1500. Single crystals were cryoprotected using 10% (v/v) glycerol mixed in with the solution that yielded the crystals and plunge-frozen in liquid nitrogen.

X-ray diffraction data were collected at the Australian Synchrotron using the Macromolecular Crystallography MX2 beamline (microfocus) at 100 K using a wavelength of 0.9537 Å (20). The auto-processed data from the Australian Synchrotron (processed using the XDS pipeline) were further processed using AIMLESS from the CCP4i suite (21,22). Initial phases were obtained using PhaserMR to perform molecular replacement with the structure of BRD3-BD1 from PDB ID: 3S91 as a molecular replacement model (23). COOT was used for manual building of the peptides (24). For all structures described in this, initial attempts to build the peptides revealed additional density that corresponded to an additional copy of the peptides rotated ∼180° around the axis perpendicular to the α-helical backbone of the peptides. The best phases were obtained by modelling the two rotated copies of the peptides into the density at 50% occupancy each. The structures were refined using iterative rounds of manual model building in COOT followed by refinement using Phenix refine (25). The data collection and refinement statistics for all structures described in this study are outlined in **Table S2**. The final models were deposited to and validated by the PDB. All structure diagrams presented in this paper are generated using PyMOL (Schrödinger). Buried surface areas were determined using the PISA server (https://www.ebi.ac.uk/pdbe/pisa/;).

## Supporting information

SupportingData

## DATA AVAILABILITY

The coordinates and structure factors for all structures described in this paper have been deposited in the Protein Data Bank (PDB). The PDB IDs for all structures described in this study are as follows: 7TO8 (BRD3-BD1 in complex with 2AcK), 7TO9 (BRD3-BD1 in complex with 2AcK.4E), 7TO7 (BRD3-BD1 in complex with 1AcK.4E). The PDB ID codes for each structure have also been provided throughout the main text and are also listed in in the Supporting Information.

## SUPPORTING INFORMATION

This article contains supporting information provided as a separate file.

## ACKNOWLEDGEMENTS

This research was undertaken using the MX1 and MX2 beamlines at the Australian Synchrotron, part of Australian Nuclear Science and Technology Organization, and made use of the Australian Cancer Research Foundation Eiger 16M detector. We acknowledge the SPR and NMR nodes of the Sydney Analytical Core Facilities The University of Sydney for providing access to SPR infrastructure.

## AUTHOR CONTRIBUTIONS

Conceptualization, L.J.W., T.P. and J.P.M.; Methodology, J.K.K.L., L.J.W., K.P., T.P., H.S. and J.P.M.; Investigation, J.K.K.L., L.J.W., K.P. and J.P.M.; Formal analysis, J.K.K.L., L.J.W., K.P., P.S., P.P., N.J., and J.P.M.; Resources, A.N., R.J.P., H.S. and J.P.M.; Writing, J.K.K.L., L.J.W., K.P. and J.P.M.; Supervision, R.J.P., T.P., H.S. and J.P.M.

## FUNDING AND ADDITIONAL INFORMATION

J.P.M., L.J.W., and R.J.P. received funding from the National Health and Medical Research Council (APP 1161623). R.J.P. is supported by an Investigator Grant (APP 1174941) from the National Health and Medical Research Council. This work was also supported by the Francis Crick Institute which receives its core funding from Cancer Research UK (CC2030), the UK Medical Research Council (CC2030), and the Wellcome Trust (CC2030).

For the purpose of Open Access, the authors have applied a CC BY public copyright licence to any Author Accepted Manuscript version arising from this submission.

