## SupportingData for "mRNA display reveals a class of high-affinity bromodomain-binding motifs that are not found in the human proteome"

### LIST OF MATERIALS INCLUDED

#### INCLUDED IN THIS DOCUMENT:

- **Supplementary Table S2**
- **Supplementary Figure S1 – S10**

#### PROVIDED AS SEPARATE FILES:

**Table S1. Enriched sequences from four rounds of RaPID screening of the 1AcK and 2AcK libraries against BRD3-BD1.**

**Table S2.** Data collection and refinement statistics for the crystal structure of BRD3-BD1 in complex with 2AcK (PDB ID: 7TO8), BRD3-BD1 in complex with 2AcK.4E (PDB ID: 7TO9), and BRD3-BD1 in complex with 1AcK.4E (PDB ID: 7TO7) solved using molecular replacement.

| | BRD3-BD1:2AcK | BRD3-BD1:2AcK.4E | BRD3-BD1:1AcK.4E |
|---|---|---|---|
| **Data collection** | | | |
| Space group | P 1 21 1 | P 1 21 1 | P 21 21 21 |
| Cell dimentions | | | |
| *a, b, c* (Å) | 40.84, 64.43, 53.47 | 40.58, 64.37, 52.31 | 41.86, 99.28, 120.61 |
| *α, β, γ* (°) | 90.00, 95.03, 90.00 | 90.00, 94.94, 90.00 | 90.00, 90.00, 90.00 |
| Resolution (Å) | 1.50 (1.50-1.53) | 1.60 (1.60-1.63) | 1.93 (1.93-1.98) |
| $R_{merge}$ | 0.044 (0.373) | 0.048 (0.673) | 0.129 (1.118) |
| $I/\sigma I$ | 20.0 (4.6) | 12.0 (1.7) | 11.3 (1.8) |
| CC(1/2) | 0.999 (0.940) | 0.999 (0.785) | 0.998 (0.636) |
| Completeness (%) | 99.3 (98.1) | 99.9 (100.0) | 99.6 (94.4) |
| Redundancy | 6.8 (7.1) | 4.4 (4.4) | 8.2 (8.1) |
| **Refinement** | | | |
| Resolution (Å) | 1.50 | 1.60 | 1.93 |
| No. reflections | 43884 | 35386 | 39319 |
| $R_{work}/R_{free}$ | 0.1770/0.2001 | 0.1985/0.2268 | 0.2013/0.2506 |
| No. atoms | 2439 | 2356 | 4478 |
| Protein | 1960 | 1959 | 3926 |
| Ligand/ion | 248 | 222 | 396 |
| Water | 231 | 175 | 156 |
| *B*-factors | 23.00 | 29.00 | 27.00 |
| R.m.s. deviations | | | |
| Bond lengths (Å) | 0.0058 | 0.0063 | 0.0072 |
| Bond angles (°) | 0.8326 | 0.8664 | 0.8541 |

*Values in the parentheses are for the highest resolution shell.
All data were collected on a single crystal.

**FIGURES**

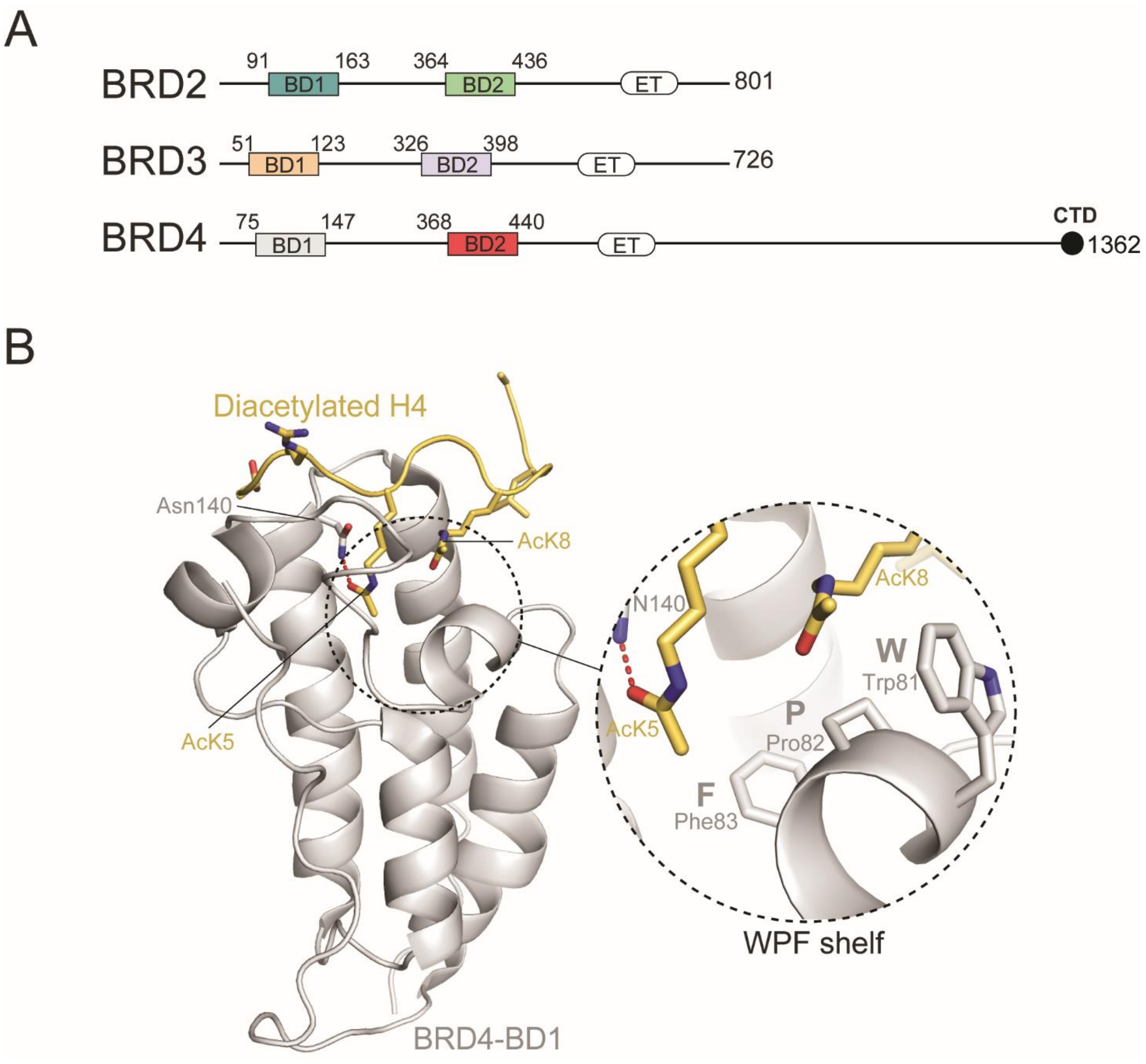


**Figure S1. The BET family of bromodomain containing proteins. A.** Topology diagrams of human BRD2, BRD3 and BRD4, showing the location of the BDs used in this study. BD stands for bromodomain, ET stands for extra-terminal domain, and CTD stands for *C*-terminal domain. **B.** The X-ray crystal structure of BRD4-BD1 (*grey*) in complex with a diacetylated Histone H4 peptide (*yellow*, PDB ID: 3UVW). *Left*: Ribbon representation of the BRD4-BD1:diacetylated Histone H4 complex. The conserved asparagine (Asn140) that makes a hydrogen bond with the N-terminal AcK residue from the diacetylated Histone H4 motif (AcK5) are displayed as sticks and the hydrogen bond formed between them is indicated by the *orange dashed line*. *Right inset:* A close-up of the WPF shelf that forms significant van der Waals interactions with C-terminal AcK (AcK8) of the diacetylated motif in Histone H4 is shown in the diagram enclosed in the *dashed circle*.

| 1st | 2nd: U | C | A | G | 3rd |
|---|---|---|---|---|---|
| U | Phe | Ser | Tyr | Cys | U |
| | | | | | C |
| | Leu | Ser | Stop | Stop | A |
| | | | | | G |
| C | Leu | Pro | His | Arg | U |
| | | | | | C |
| | Leu | Pro | Gln | Arg | A |
| | | | | | G |
| A | Ile | Thr | Asn | Ser | U |
| | | | | | C |
| | Ile | Thr | Lys | Arg | A |
| | KAc | | | | G |
| G | Val | Ala | Asp | Gly | U |
| | | | | | C |
| | Val | Ala | Glu | Gly | A |
| | | | | | G |

Initiator = Ac-Ala

No Trp or Met in random region

**Figure S2**. **Codon table for the RaPID screens showing amino acid assignment for each codon encoded in the randomised library.** Methionine was replaced by AcK and the initiator tRNA charged with $^{N}$Ac-Ala.

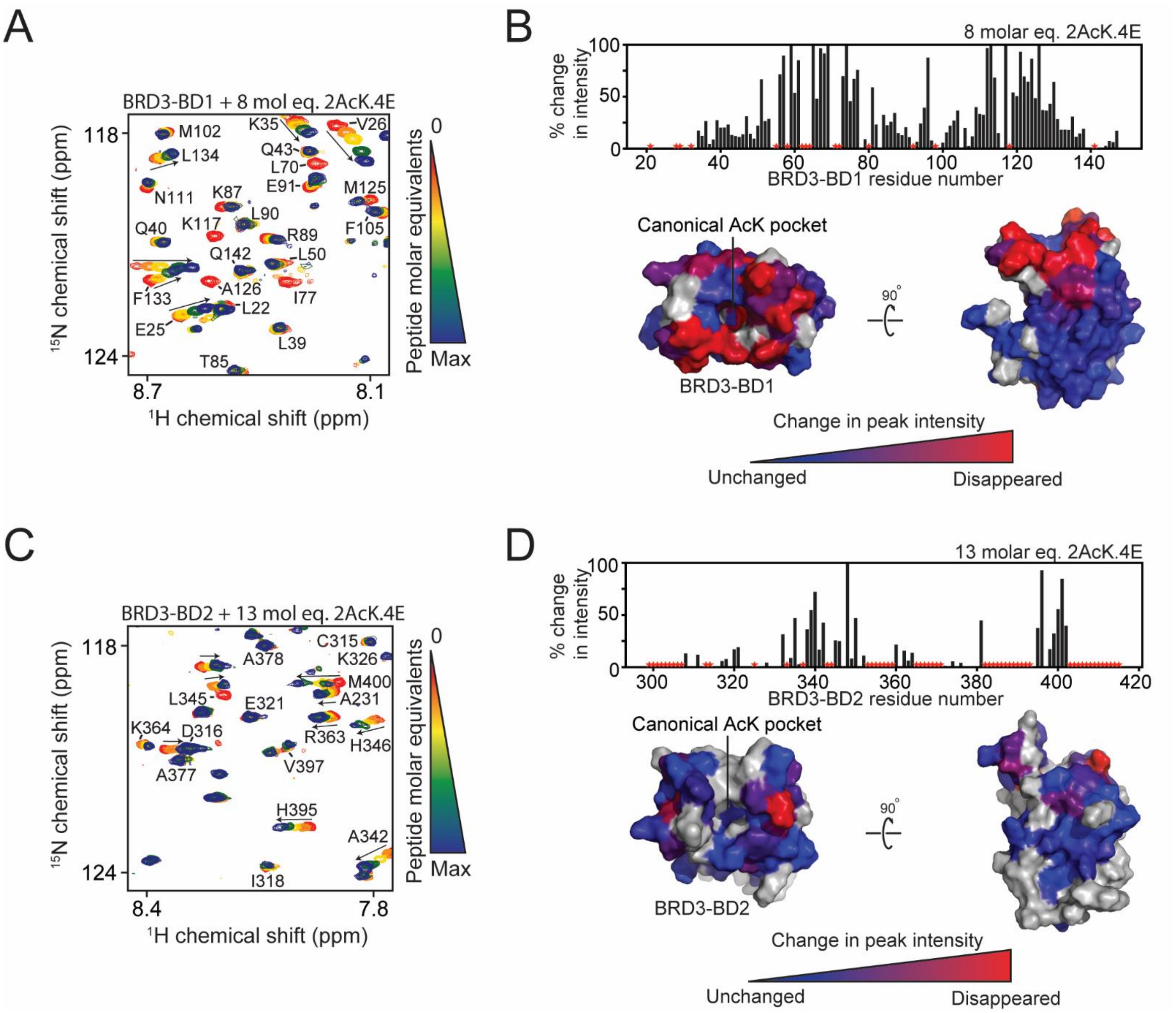


**Figure S3. $^{15}$N-HSQC titrations for 2AcK.4E into the BDs of BRD3 and structural mapping of the interactions.** For all $^{15}$N-HSQC titrations, BDs were used at a concentration of ~30 µM and the assignments and direction of movement of some signals are indicated. Residues that could not be assigned in the HSQCs are indicated by red stars in the graphs plotting intensity change and in grey on the structure of BDs. **A.** $^{15}$N-HSQC spectra of BRD3-BD1 alone (*red*) and in the presence of increasing concentrations of 2AcK.4E (up to 8 molar equivalents of the peptide, *blue*). **B.** *Top:* Quantitation of change in $^{15}$N-HSQC signal intensity for each signal following the addition of the indicated amount of 2AcK.4E to BRD3-BD1. *Bottom:* Intensity changes from the graph above mapped onto the structure of BRD3-BD1. The magnitude of the change is represented by a *blue–red* colour gradient (*red* indicates the largest change). **C.** $^{15}$N-HSQC spectra of BRD3-BD2 alone (*red*) and in the presence of increasing concentrations of 2AcK.4E (up to 13 molar equivalents of the peptide, *blue*). **D.** *Top:* Quantitation of change in $^{15}$N-HSQC signal intensity for each signal following the addition of the indicated amount of 2AcK.4E to BRD3-BD2. *Bottom:* Intensity changes from the graph above mapped onto the structure of BRD3-BD2. The magnitude of the change is represented by a *blue–red* colour gradient (*red* indicates the largest change).

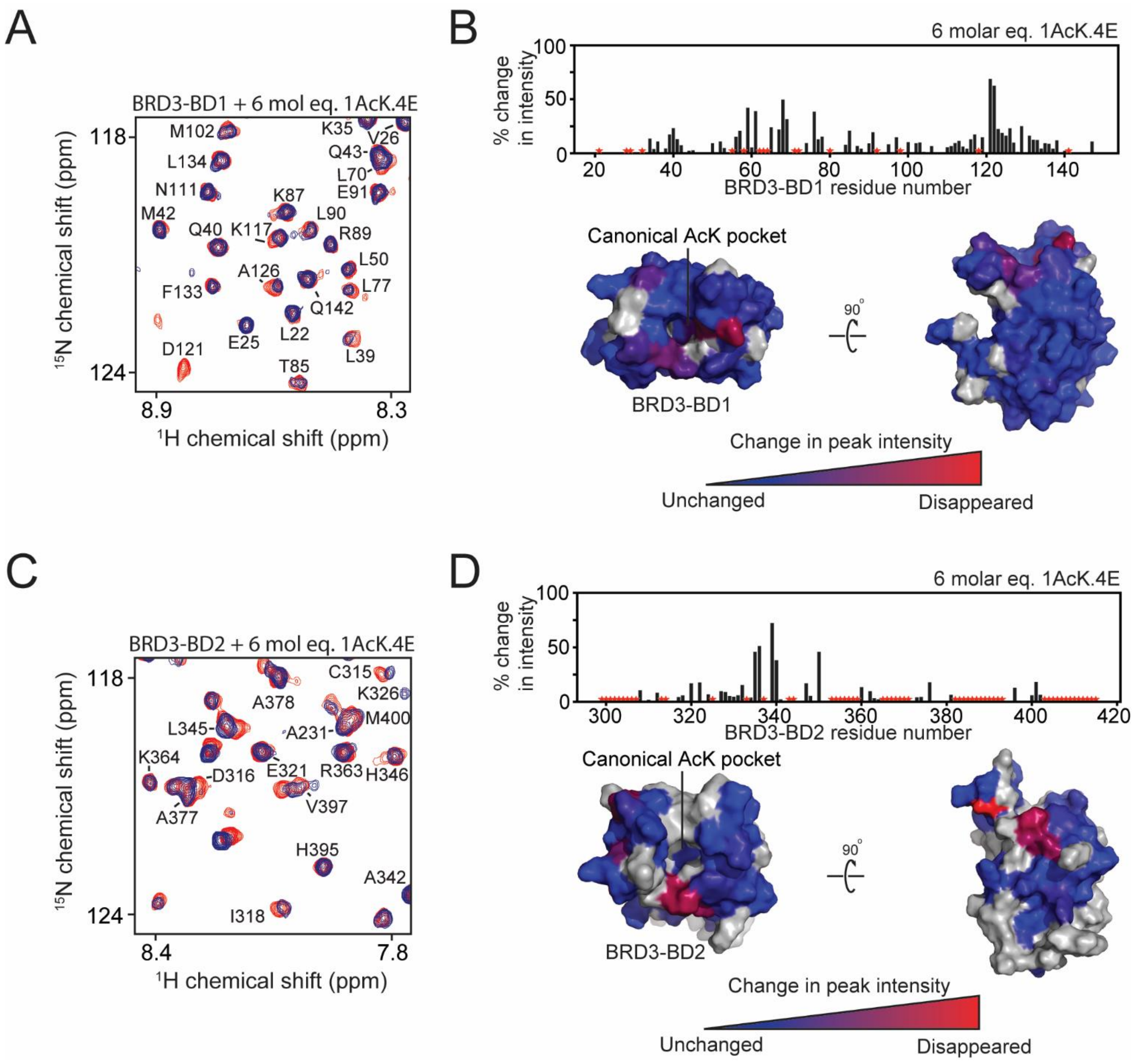


**Figure S4. $^{15}$N-HSQC titrations for 1AcK.4E into the BDs of BRD3 and structural mapping of the interactions.** For all $^{15}$N-HSQC titrations, BDs were used at a concentration of ~30 µM and the assignments and direction of movement of some signals are indicated. Residues that could not be assigned in the HSQCs are indicated by red stars in the graphs plotting intensity change and in grey on the structure of BDs. **A.** $^{15}$N-HSQC spectra of BRD3-BD1 alone (*red*) and in the presence of 6 molar equivalents of 1AcK.4E (*blue*). **B.** *Top:* Quantitation of change in $^{15}$N-HSQC signal intensity for each signal following the addition of the indicated amount of 1AcK.4E to BRD3-BD1. *Bottom:* Intensity changes from the graph above mapped onto the structure of BRD3-BD1. The magnitude of the change is represented by a *blue–red* colour gradient (*red* indicates the largest change). **C.** $^{15}$N-HSQC spectra of BRD3-BD2 alone (*red*) and in the presence of 6 molar equivalents of 1AcK.4E (*blue*). **D.** *Top:* Quantitation of change in $^{15}$N-HSQC signal intensity for each signal following the addition of the indicated amount of 1AcK.4E to BRD3-BD2. *Bottom:* Intensity changes from the graph above mapped onto the structure of BRD3-BD2. The magnitude of the change is represented by a *blue–red* colour gradient (*red* indicates the largest change).

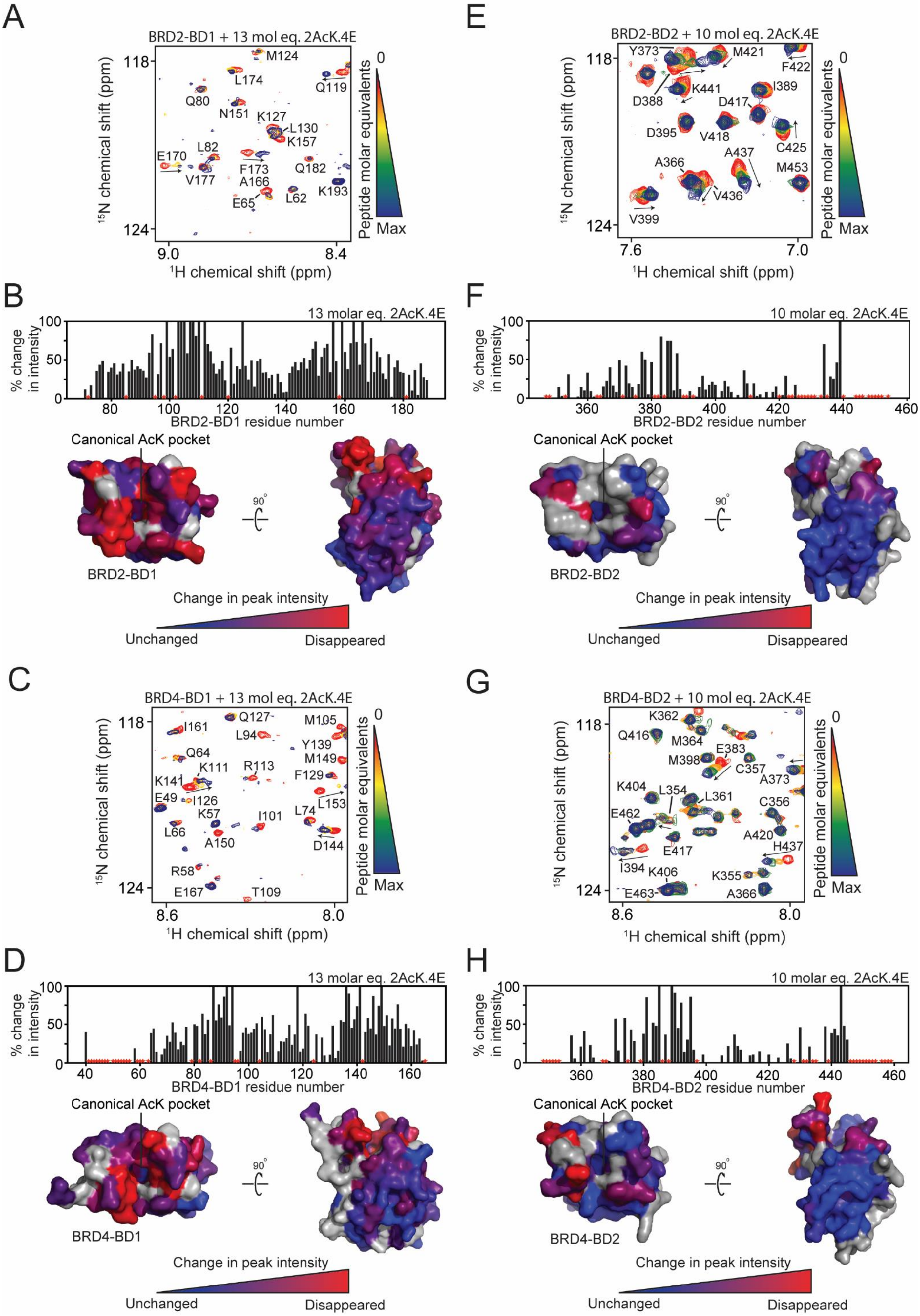
A
BRD2-BD1 + 13 mol eq. 2AcK.4E
M124
L174
Q119
Q80
N151
K127
L130
K157
L82
E170
V177
F173
Q182
A166
K193
E65
L62
$^{15}$N chemical shift (ppm)
$^{1}$H chemical shift (ppm)
118
124
9.0
8.4
Peptide molar equivalents
0
Max
B
13 molar eq. 2AcK.4E
% change in intensity
BRD2-BD1 residue number
Canonical AcK pocket
BRD2-BD1
90°
Change in peak intensity
Unchanged
Disappeared
C
BRD4-BD1 + 13 mol eq. 2AcK.4E
I161
Q127
M105
L94
Y139
Q64
M149
K141
K111
R113
F129
E49
I126
L153
K57
I101
L74
L66
A150
D144
R58
E167
T109
8.6
8.0
D
BRD4-BD1 residue number
BRD4-BD1
E
BRD2-BD2 + 10 mol eq. 2AcK.4E
Y373
M421
F422
D388
K441
I389
D417
D395
V418
C425
A437
A366
M453
V436
V399
7.6
7.0
F
10 molar eq. 2AcK.4E
BRD2-BD2 residue number
BRD2-BD2
G
BRD4-BD2 + 10 mol eq. 2AcK.4E
K362
Q416
M364
M398
E383
C357
A373
K404
L354
L361
C356
E462
E417
A420
H437
I394
K406
K355
E463
A366
H
BRD4-BD2 residue number
BRD4-BD2

**Figure S5. Peptide 2AcK.4E binds the BDs from BRD2 and BRD4 at the canonical binding site.** For all $^{15}$N-HSQC titrations, BDs were used at a concentration of ~30 µM and the assignments and direction of movement of some signals are indicated. Residues that could not be assigned in the HSQCs are indicated by red stars in the graphs plotting intensity change and in grey on the structure of BDs. **A.** $^{15}$N-HSQC spectra of BRD2-BD1 alone (*red*) and in the presence of increasing concentrations of 2AcK.4E (up to 13 molar equivalents of the peptide, *blue*). **B.** *Top:* Quantitation of change in $^{15}$N-HSQC signal intensity for each signal following the addition of the indicated amount of 2AcK.4E to BRD2-BD1. *Bottom:* Intensity changes from the graph above mapped onto the structure of BRD2-BD1. The magnitude of the change is represented by a *blue–red* colour gradient (*red* indicates the largest change). **C.** $^{15}$N-HSQC spectra of BRD4-BD1 alone (*red*) and in the presence of increasing concentrations of 2AcK.4E (up to 13 molar equivalents of the peptide, *blue*). **D.** *Top:* Quantitation of change in $^{15}$N-HSQC signal intensity for each signal following the addition of the indicated amount of 2AcK.4E to BRD4-BD1. *Bottom:* Intensity changes from the graph above mapped onto the structure of BRD4-BD1. The magnitude of the change is represented by a *blue–red* colour gradient (*red* indicates the largest change). **E.** $^{15}$N-HSQC spectra of BRD2-BD2 alone (*red*) and in the presence of increasing concentrations of 2AcK.4E (up to 10 molar equivalents of the peptide, *blue*). **F.** *Top:* Quantitation of change in $^{15}$N-HSQC signal intensity for each signal following the addition of the indicated amount of 2AcK.4E to BRD2-BD2. *Bottom:* Intensity changes from the graph above mapped onto the structure of BRD2-BD2. The magnitude of the change is represented by a *blue–red* colour gradient (*red* indicates the largest change). **G.** $^{15}$N-HSQC spectra of BRD4-BD2 alone (*red*) and in the presence of increasing concentrations of 2AcK.4E (up to 10 molar equivalents of the peptide, *blue*). **H.** *Top:* Quantitation of change in $^{15}$N-HSQC signal intensity for each signal following the addition of the indicated amount of 2AcK.4E to BRD4-BD2. *Bottom:* Intensity changes from the graph above mapped onto the structure of BRD4-BD2. The magnitude of the change is represented by a *blue–red* colour gradient (*red* indicates the largest change).

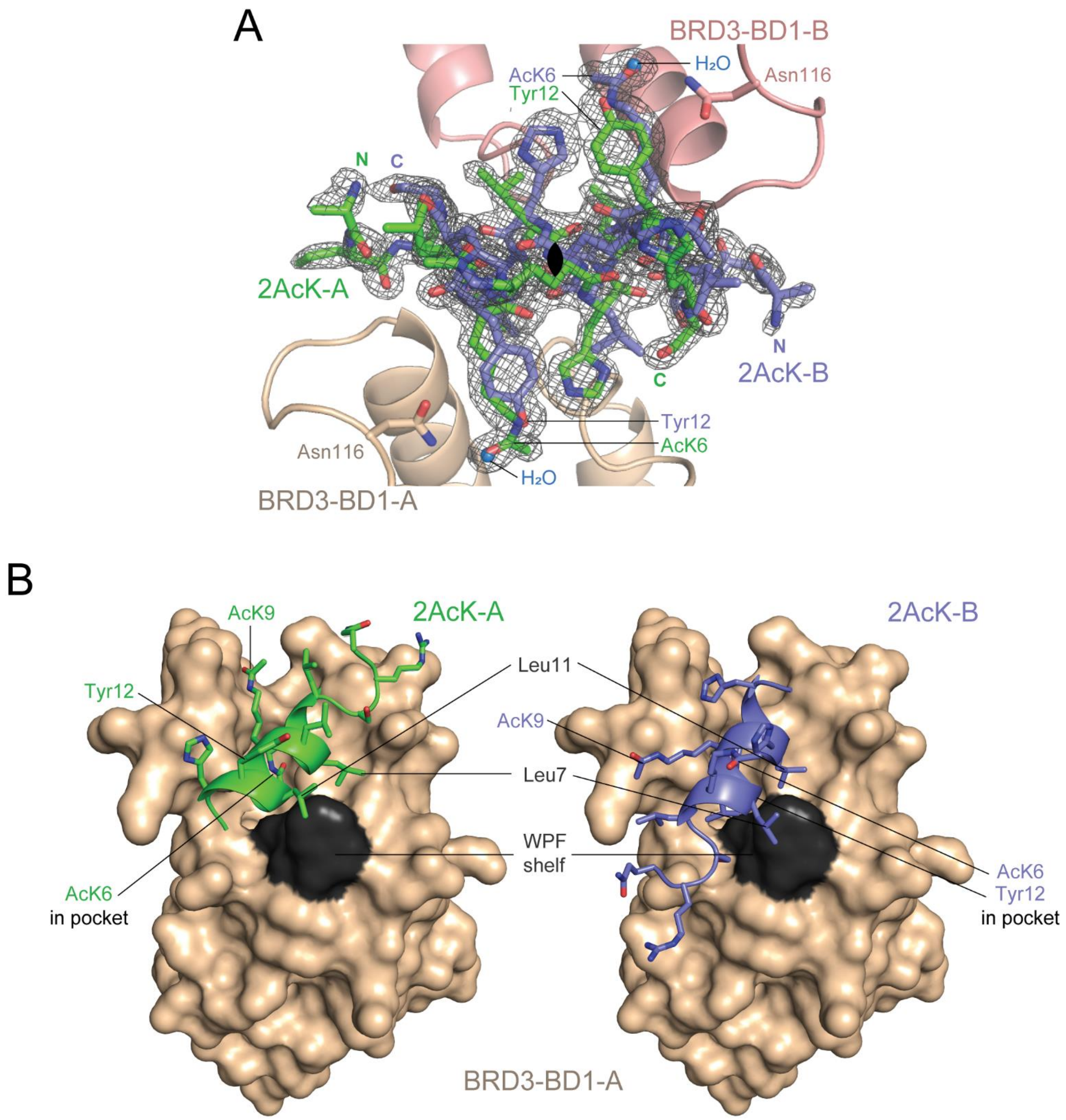


**Figure S6. Structure of BRD3-BD1 bound to 2AcK. A.** Electron density ($2F_O - F_C$) observed between the two copies of BRD3-BD1 in the asymmetric unit. The density (shown as a mesh) is clearly better fitted by using a model that incorporates two copies of the peptide (each at 50% occupancy, *green* and *purple*) rotated 180° about the indicated axis (*black ellipse*). The pseudo-two-fold axis of rotation, which maps each BD onto the position of the other and approximately maps the overall helical shape of the peptide backbone onto itself (but running in the opposite direction) is indicated with a *black ellipse*. It can be seen that AcK6 and Tyr12 of 2AcK occupy the AcK-binding pocket of BRD3-BD1-A and BRD3-BD1-B, respectively. **B.** Comparison of peptide binding geometry for binding to the two BDs in the asymmetric unit. The orientation of peptide 2AcK-B that binds BRD3-BD1-B using Tyr12 is superimposed onto BRD3-BD1-A to demonstrate the relative orientations of 2AcK.

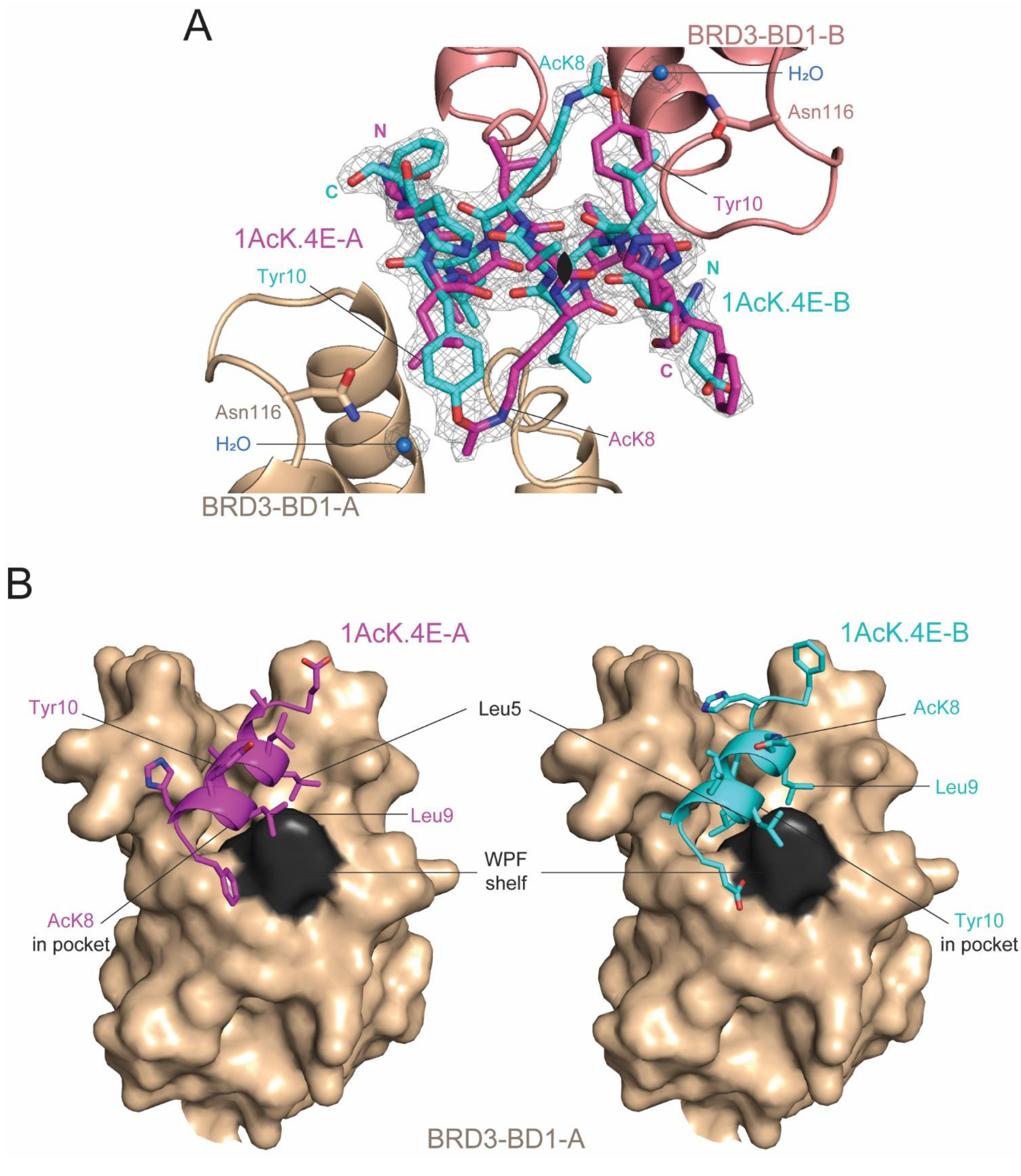


**Figure S7. Structure of BRD3-BD1 bound to 1AcK.4E. A.** Electron density ($2F_0 - F_c$) observed in between the two copies of BRD3-BD1 in the asymmetric unit. The density (shown as a mesh) is clearly better fitted by using a model that incorporates two copies of the peptide (each at 50% occupancy, *magenta* and *cyan*) rotated 180° about the indicated axis (*black ellipse*). The pseudo-two-fold axis of rotation, which maps each BD onto the position of the other and approximately maps the overall helical shape of the peptide backbone onto itself (but running in the opposite direction) is indicated with a *black ellipse*. It can be seen that AcK8 and Tyr10 of 1AcK.4E occupy the AcK-binding pocket of BRD3-BD1-A and BRD3-BD1-B, respectively. **B.** Comparison of peptide binding geometry for binding to the two BDs in the asymmetric unit. The orientation of peptide 1AcK.4E-B that binds BRD3-BD1-B using Tyr10 is superimposed onto BRD3-BD1-A to demonstrate the relative orientations of 1AcK.4E.

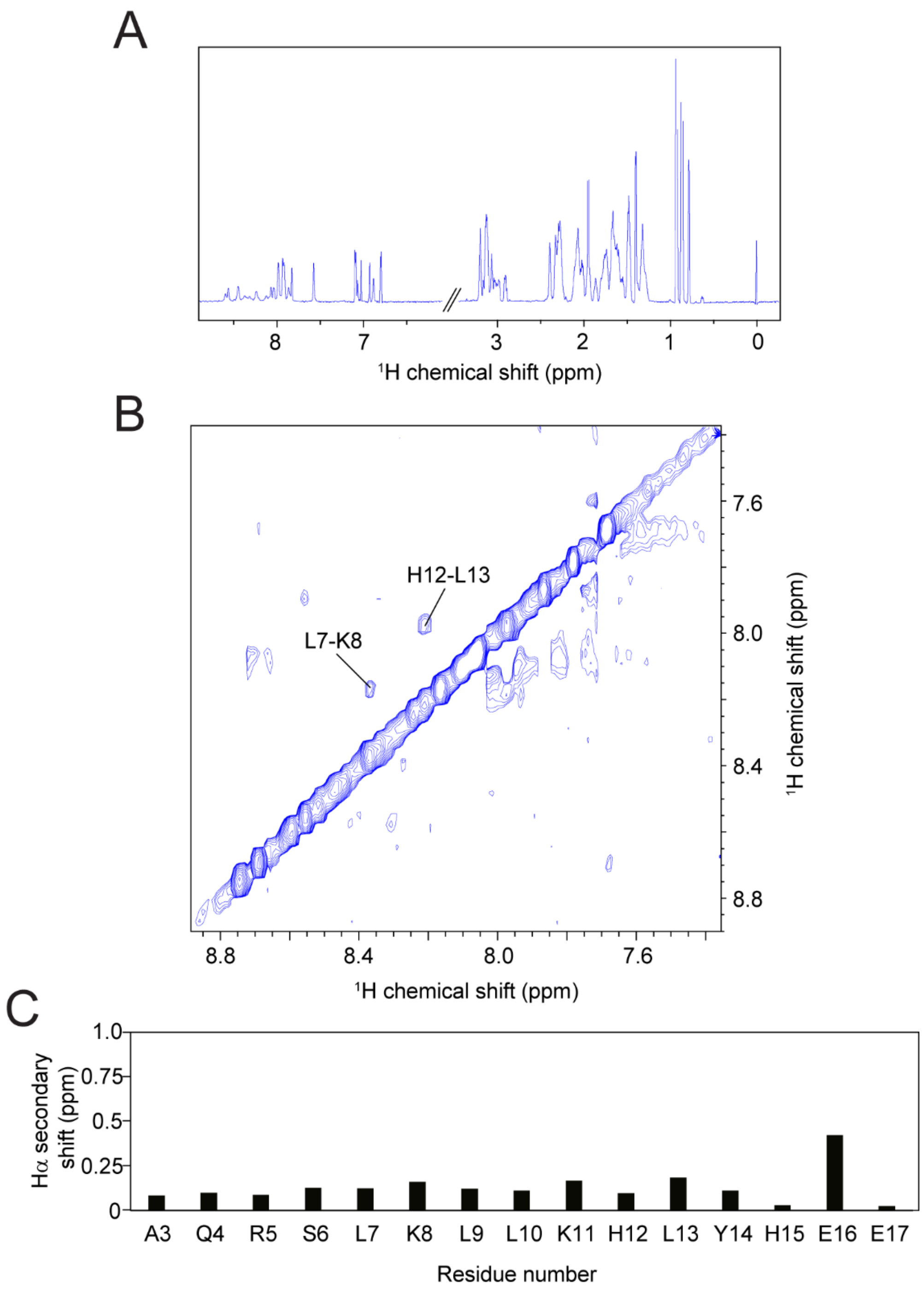


**Figure S8. 2AcK.4E is intrinsically disordered. A.** Portions of a 1D 1H NMR spectrum of 2AcK.4E (recorded at 800 MHz, 25 °C). **B**. HN-HN region of a 2D NOESY spectrum of 2AcK.4E (mixing time = 100 ms). The lack of a significant number of NOEs connecting pairs of HN protons strongly suggests that the peptide does not form appreciable amounts of α-helix. **C**. Secondary chemical shifts for Hα protons of 2AcK.4E, obtained by subtracting the random coil chemical shift for each residue from the observed chemical shift. Calculations were performed on the CSI 3.0 server of the Wishart lab (http://csi3.wishartlab.com/). The small and uniformly positive nature of the secondary shifts is consistent with a largely disordered polypeptide with a weak preference for an α-helical conformation.

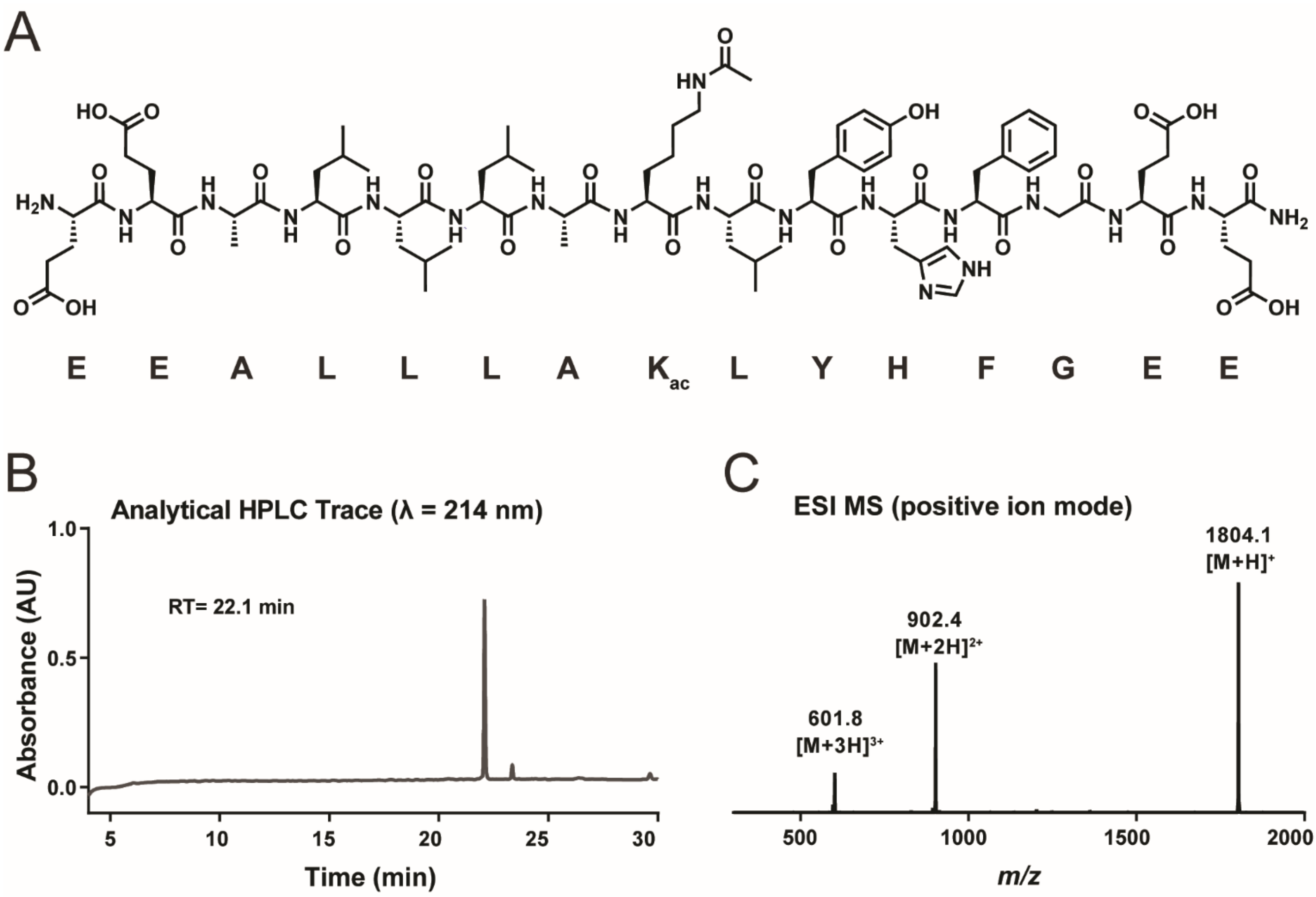


**Figure S9. HPLC trace and MS spectrum for peptide 1AcK.4E. A.** Structure and sequence of peptide 1AcK.4E. **B.** Analytical HPLC chromatogram. $R_t$ = 22.1 min (1 to 60 vol.% MeCN in $H_2O$ with 0.1 vol.% TFA over 30 min, λ = 214 nm). **C.** Low resolution peptide mass spectrum in positive ion ESI mode. Observed masses at *m/z* 1804.1 $[M+H]^+$, 902.4 $[M+2H]^{2+}$, 601.8 $[M+3H]^{3+}$.

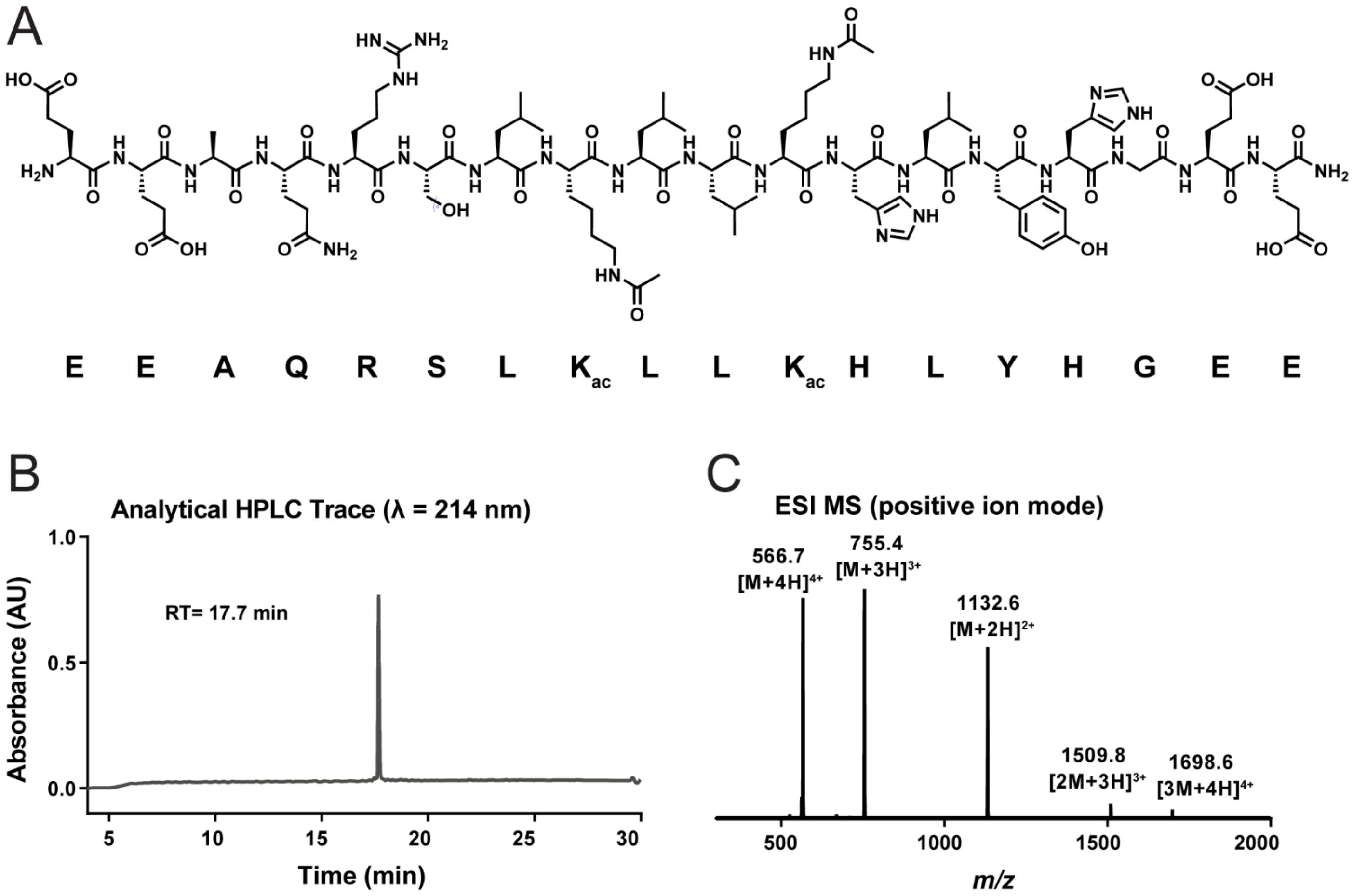


**Figure S10. HPLC trace and MS spectrum for peptide 2AcK.4E. A.** Structure and sequence of peptide 2AcK.4E. **B.** Analytical HPLC chromatogram. $R_t$ = 17.7 min (1 to 60 vol.% MeCN in $H_2O$ with 0.1 vol.% TFA over 30 min, λ = 214 nm). **C.** Low resolution peptide mass spectrum in positive ion ESI mode. Observed masses at *m/z* 1698.6 $[3M+4H]^{4+}$, 1509.8 $[2M+3H]^{3+}$, 1132.6 $[M+2H]^{2+}$, 755.4 $[M+3H]^{3+}$, 566.7 $[M+4H]^{4+}$.
